# Rapid weed adaptation and range expansion in response to agriculture over the last two centuries

**DOI:** 10.1101/2022.02.25.482047

**Authors:** Julia M. Kreiner, Sergio M. Latorre, Hernán A. Burbano, John R. Stinchcombe, Sarah P. Otto, Detlef Weigel, Stephen I. Wright

## Abstract

North America has seen a massive increase in cropland use since 1800, accompanied more recently by the intensification of agricultural practices. Through genome analysis of present-day and historical samples spanning environments over the last two centuries, we studied the impact of these changes in farming on the extent and tempo of evolution across the native range of common waterhemp (*Amaranthus tuberculatus)*, a now pervasive agricultural weed. Modern agriculture has imposed strengths of selection rarely observed in the wild, with striking shifts in allele frequency trajectories since agricultural intensification in the 1960s. An evolutionary response to this extreme selection was facilitated by a concurrent human-mediated range shift. By reshaping genome-wide diversity across the landscape, agriculture has driven the success of this weed in the 21^st^-century.

**One Sentence Summary:** Modern agriculture has shaped the evolution of a native plant into a weed by driving range shifts and strengths of selection rarely observed in the wild.

## Main text

Agricultural practices across North America have rapidly intensified over the last two centuries, through cropland expansion (*1*), monoculture plantings (*2, 3*), and increased chemical inputs (*4, 5*). Since the beginning of the 1800s, cropland usage has expanded from 8 million to 200 million hectares in Canada and the United States alone (*1*). Since the mid 1900’s, development of new crop varieties–including high-yield and herbicide-resistant wheat, corn, and soy (*6, 7*)—have greatly improved the efficiency of food production in all farming sectors. Combined with increased reliance on pesticides, fertilizers, irrigation, and large-scale mechanization, this global transformation is oft-referenced as the agricultural “Green Revolution” (*8–10*). Pesticide effectiveness, however, has been limited by the evolution of resistance across numerous pest species (*11–14*). While technological innovation for efficient food production has risen with increasing global food demands, the concomitant landscape conversion has become one of the foremost drivers of global biodiversity loss (*15*).

Species that have managed to survive, and even thrive, in the face of such extreme environmental change provide remarkable examples of rapid adaptation on contemporary timescales and illustrate the evolutionary consequences of anthropogenic change. One such species is common waterhemp (*Amaranthus tuberculatus*), an agricultural weed that is native to North America and persists in large part in natural, riparian habitats (*16, 17*), providing a unique opportunity to investigate the timescale and extent of contemporary agricultural adaptation. The genetic changes underlying weediness are particularly important to understand in *A. tuberculatus*, as it has recently become one of the most problematic agricultural weeds in North America due to its widespread adaptation to herbicides, persistence in fields across seasons, and strong ability to compete with both soy and corn (*18, 19*). Determining the role of newly arisen mutations, genetic variants predating the onset of environmental change (*20, 21*), migration across the range (*22*), and their interactions (e.g. (*23, 24*)), will inform on the temporal and spatial scales at which contemporary adaptation occurs and management strategies should be employed.

To understand how changing agricultural practices have shaped the success of a ubiquitous weed, we analyze genomic data from contemporary paired natural and agricultural populations alongside historical herbarium samples collected from 1828 until 2011 (**Fig 1)**. With this design, we identify agriculturally adaptive alleles (i.e., those that occur at consistently higher frequencies in agricultural than in nearby natural sites), track their frequencies across nearly two centuries, and link the tempo of weed adaptation to demographic changes and key cultural shifts in modern agriculture.

**Fig 1.**
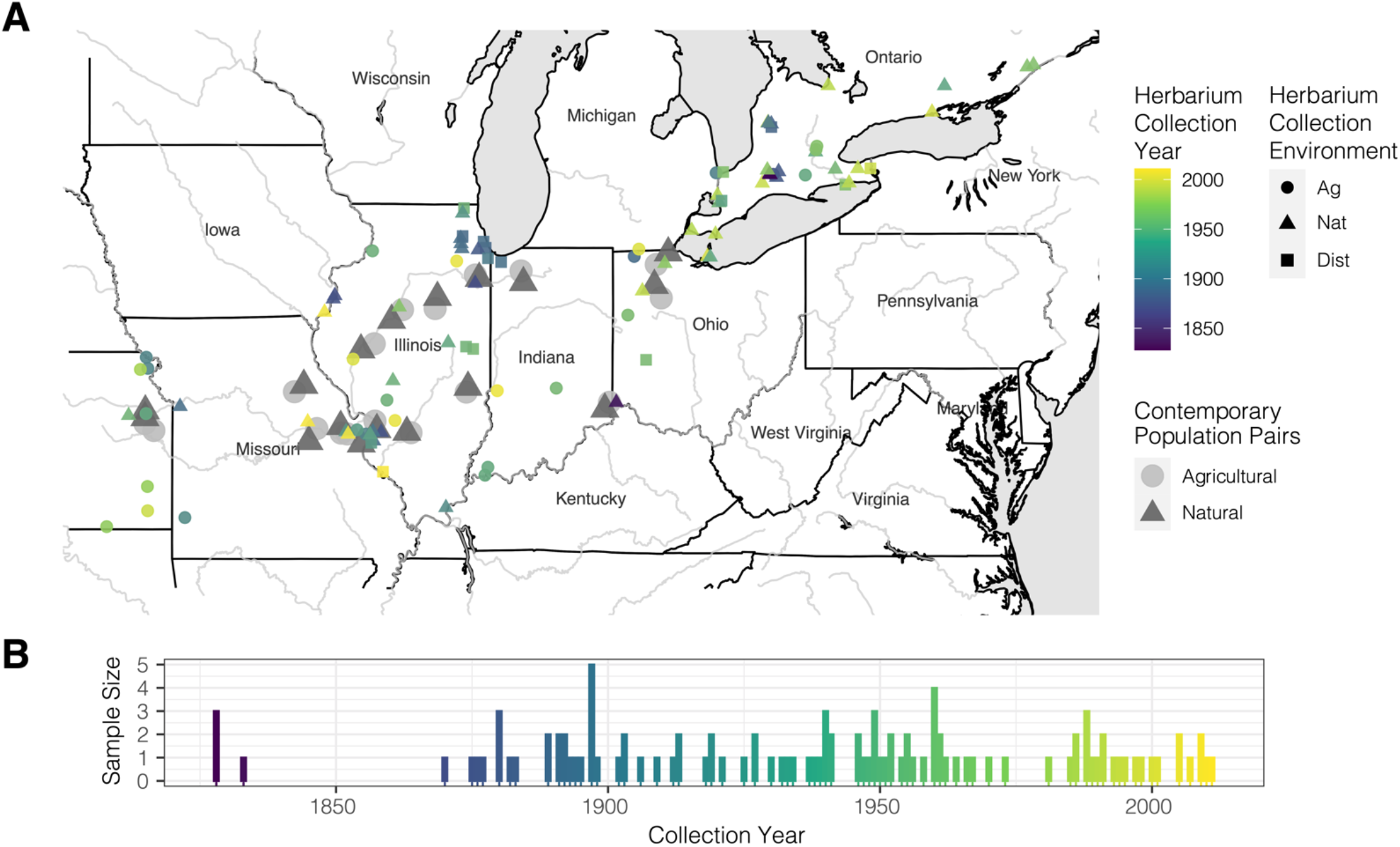
Sequenced waterhemp collections through space and time. **A)** Map of 17 contemporary paired natural-agricultural populations [n=187, collected and sequenced in Kreiner et al., 2021 (*25*)], along with 108 novel sequenced herbarium specimens dating back to 1828 collected across three environment types (Ag=Agricultural, Nat=Natural, Dist=Disturbed; metadata provided in Data S2). **B)** Distribution of sequenced herbarium samples through time.

### The genome-wide signatures of agricultural adaptation

To find alleles favored under current farming practices, we looked for those that were consistently overrepresented in extant populations collected in agricultural habitats compared to neighboring riparian (“natural”) habitats (*25*) using Cochran–Mantel– Haenszel (CMH) tests (**Fig 2A**). Alleles associated with agricultural environments (the 0.1% of SNPs with lowest CMH p-values; n=7264) are significantly enriched for 29 GO-biological process terms related to growth and development, reproduction, cellular metabolic processes, and responses to abiotic, endogenous and external stimuli, including response to chemicals (**Table S1**). The importance of chemical inputs in shaping weed agricultural adaptation is clear in that the most significant agriculturally associated SNP (raw p-value = 8.551×10^−11^, [FDR corrected] q-value = 0.00062) falls just 80 kb outside the gene protoporphyrinogen oxidase (*PPO*)—the target of PPO-inhibiting herbicides (**Fig 2B**). PPO herbicides were widely used in the 1990s, but have seen a recent resurgence to control and slow the spread of glyphosate resistant weeds (*26, 27*). Other genes with the strongest agricultural associations include *ACO1*, which has been shown to confer oxidative stress tolerance (*28*); *HB13*, involved in pollen viability (*29*) as well as drought and salt tolerance (*30*); *PME3*, involved in growth via germination timing (*31*); *CAM1*, a regulator of senescence in response to stress (*32, 33*); and both *CRY2* and *CPD*, two key regulators of photomorphogenesis and flowering via brassinosteroid signaling (*34–37*) (**Table S2)**. Natural-vs-agricultural F_ST_ (allele frequency differentiation) is highly correlated with the CMH test statistic (Pearson’s *r* = 0.987), with 78% [98%] of CMH focal SNPs overlapping with the top 0.01% [0.1%] of F_ST_ hits (**Fig S1)**. Despite negligible genome-wide differentiation among environments suggesting widespread gene flow (F_ST_ = 0.0008; with even lower mean F_ST_ between paired sites = -0.0029; **Fig 2C)**, our results suggest that strong antagonistic selection acts to maintain spatial differentiation for particular alleles—403 SNPs with a CMH q-value < 0.10.

**Fig 2.**
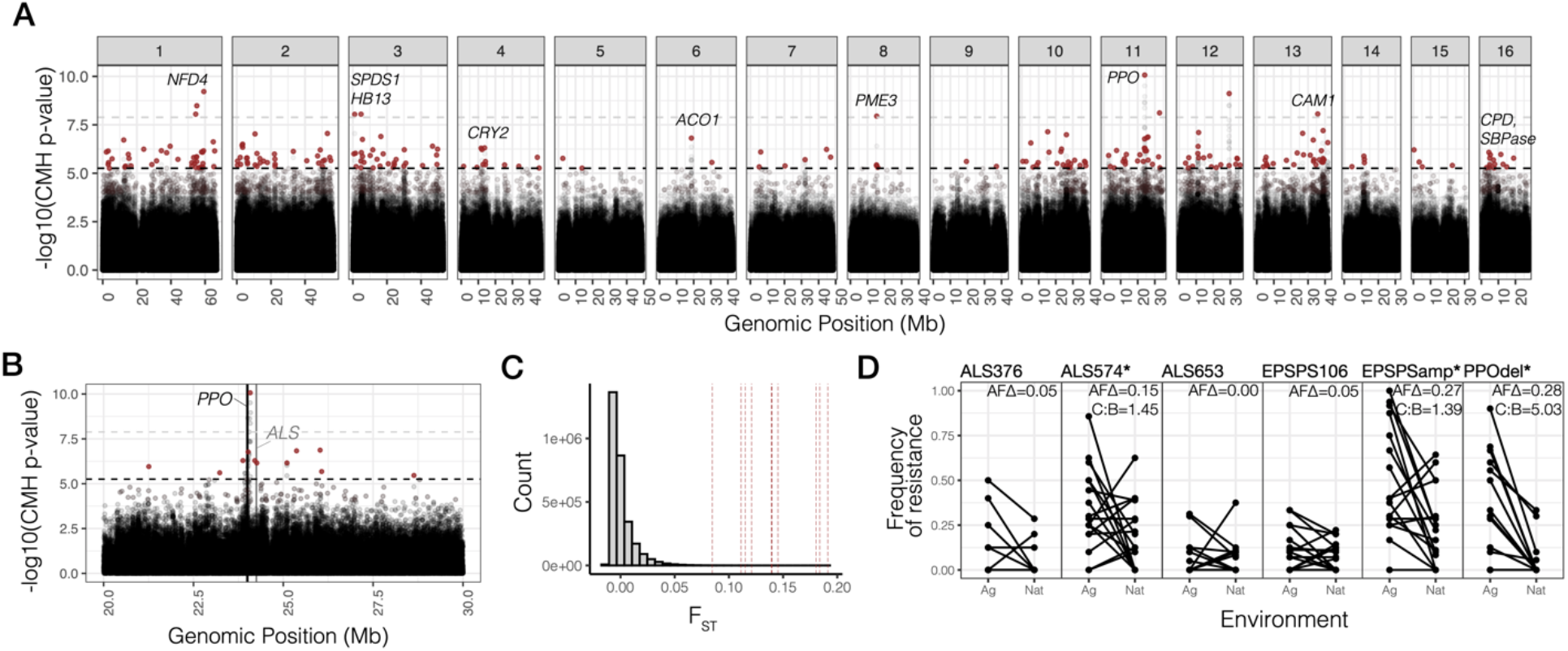
Signals of contemporary agricultural adaptation, gene flow, and antagonistic selection across the genome in *A. tuberculatus*. **A)** Results from Cochran–Mantel–Haenszel (CMH) tests for SNPs with consistent differentiation among environments across contemporary natural-agricultural population pairs. A 10% FDR threshold is indicated by the lower dashed horizontal black line, while the Bonferroni corrected p-value < 0.1 cut-off is shown by the upper dashed horizontal gray line. Red points indicate focal agricultural-associated SNPs after aggregating linked variation (r^2^ > 0.25 within 1 Mb). Candidate agriculturally adaptive genes for peaks that are significant at a 10% FDR threshold shown. **B)** CMH results from the scaffold containing the most significant CMH p-value, corresponding to variants linked to the PPO210 deletion conferring herbicide resistance and to the nearby herbicide-targeted gene *ALS*. **C)** Distribution of F_ST_ values between all agricultural and natural samples for ∼3 million genome-wide SNPs (minor allele frequency > 0.05). Vertical lines indicate F_ST_ values for the 10 candidate genes named in A. **D)** Population-level frequencies of six common herbicide resistance alleles across geographically paired agricultural and natural habitats sampled in 2018 (pairs connected by horizontal lines). The first four columns are nonsynonymous variants in *ALS* and *EPSPS*, followed by EPSPSamp (a 10 Mb-scale amplification that includes *EPSPS*), and an in-frame single-codon deletion in PPO. Estimates of per-migrant natural cost: agricultural benefit ratio (C:B) is shown in the top right corner, for the three resistance alleles with significant (*) allele frequency differences (AFΔ) across environment types in a linear regression.

To further investigate the extent to which herbicides shape adaptation to agriculture, we assayed patterns of environmental differentiation at known resistance variants. Eight such alleles were present in contemporary samples, only six of which were common **(Table S3)**: a deletion of codon 210 within *PPO* (*38*), a copy number amplification and a non-synonymous mutation within 5-enolpyruvylshikimate-3-phosphate synthase (*EPSPS*) conferring resistance to glyphosate herbicides (*39*), and 3 separate non-synonymous mutations within acetolactate synthase (*ALS*) conferring resistance to ALS-inhibiting herbicides (*19*). While these resistance alleles were at intermediate frequency in agricultural populations, ranging from 0.08 to 0.35, they tended to be rarer but still frequent in natural populations, ranging from 0.04 to 0.22 (**Fig 2C)**. Three out of six common resistance alleles show significant allele frequency differences among environments (EPSPSamp: *F =* 8.74, *p* = 0.006; PPO210: *F =* 40.98; *p* = 1.25e-09; ALS574: *F =* 6.28; *p* = 0.013), two of which represent among the strongest signals of differentiation genome-wide. Natural-vs-agricultural F_ST_ at the PPO210 deletion, 0.21, is higher than anywhere else in the genome and is even stronger when calculated within population pairs (F_ST_= 0.27) (**Fig 2C**). Similarly, the EPSPS amplification is ranked 20th among genome-wide biallelic F_ST_ values, 0.14 (within-pair F_ST_= 0.22), in support of herbicides as a foremost driver of agricultural adaptation (**Fig 2D)**.

To infer the importance of selective trade-offs in adaptation across natural and agricultural environments, we implemented a Wright-Fisher allele-frequency-based migration-selection balance model for these three differentiated resistance alleles, as well as the top 30 independent CMH outliers. Assuming these alleles are at a steady-state between migration and selection, we inferred that the costs of resistance per migrant that has arrived into natural environments are consistently higher than the benefits of resistance per migrant that has arrived into agricultural environments (per-migrant cost: benefit ratio ranges from 1.39 for EPSPSamp and 1.45 for ALS574, to 5.03 for the PPO210 deletion; **Fig 2D, Table S3**). Thus, the spread of these three common herbicide resistance alleles appears to be constrained either by more consistent selection against resistance in herbicide-free, natural environments, or by particularly high rates of migration of susceptible alleles from natural into agricultural environments. In comparison, for the top 30 independent CMH outliers, the costs per migrant that has arrived in natural environments were about equally likely to be stronger or weaker (12/28, 42%) than the benefits per migrant in agricultural environments (**Fig S2)**. This approach provides a novel and sensitive alternative to experimental studies of fitness costs which vary greatly depending on context (*40*), highlighting the potentially important role of resistance costs across a diverse set of individuals within actual agronomic and natural environments. In these field settings, further work is necessary to understand the contributions of temporal and spatial heterogeneity in both migration and selection for and against resistance across the landscape.

### Agriculturally-adaptive alleles change rapidly with intensified regimes

With a genome-wide set of agriculture-associated alleles (251 loci after aggregating linked SNPs), we searched for signatures of temporal evolution using newly collected whole genome sequence data from a set of historical herbarium samples (n=108) dating back to 1828. These samples provide snapshots of the genetic changes that have occurred over this time period and across environment types, with collections from natural and weedy (agricultural and disturbed) habitats **(Fig 1)**. Of the 165 loci for which we had sufficient information in the historical SNP set (sequenced to 10x coverage on average), 151 were segregating with the same reference/alternate allele combination (i.e. 11 were dropped due to multi-allelism), and only three were invariant. To model allele frequency change through time at these alleles, we implemented logistic regressions of genotypes (within individual allele frequencies) at each locus on collection year, where 2*slope of the logit-transform is equivalent to the strength of selection (*s*) in a diploid model of selection (where *s* is the fitness difference between homozygotes, assuming additivity; *see Methods* for model and simulations (*41*)).

Consistent with the rapid change in land use and farming practices in the recent past, the frequency of these 154 contemporary agricultural alleles has increased substantially over the last two centuries. Whereas in natural environments agriculturally-associated alleles have increased by 6% on average since 1870, the earliest time point at which we have collections across environment types, these same alleles have increased by 22% in disturbed and agricultural environments (**Fig 3A**). This observed change greatly exceeds the expected change over this time period, based on genome-wide patterns that reflect drift, migration, selection, and demographic change (null 95% interquantile range for allele frequency change in natural sites = [-2.7, 2.0.%]; for change in agricultural and disturbed sites = [3.3, 7.9%]). We generated these null expectations by randomly sampling a set of 154 loci with the same distribution of contemporary allele frequencies (**Fig S4**) and calculating their frequency change through time across herbarium samples, separately in each environment, 1000 times (see *Methods* (*41*)). That the observed change in natural environments is also more extreme than what is expected is consistent with ongoing migration of agriculturally-selected alleles and subsequent costs in natural environments.

**Fig 3.**
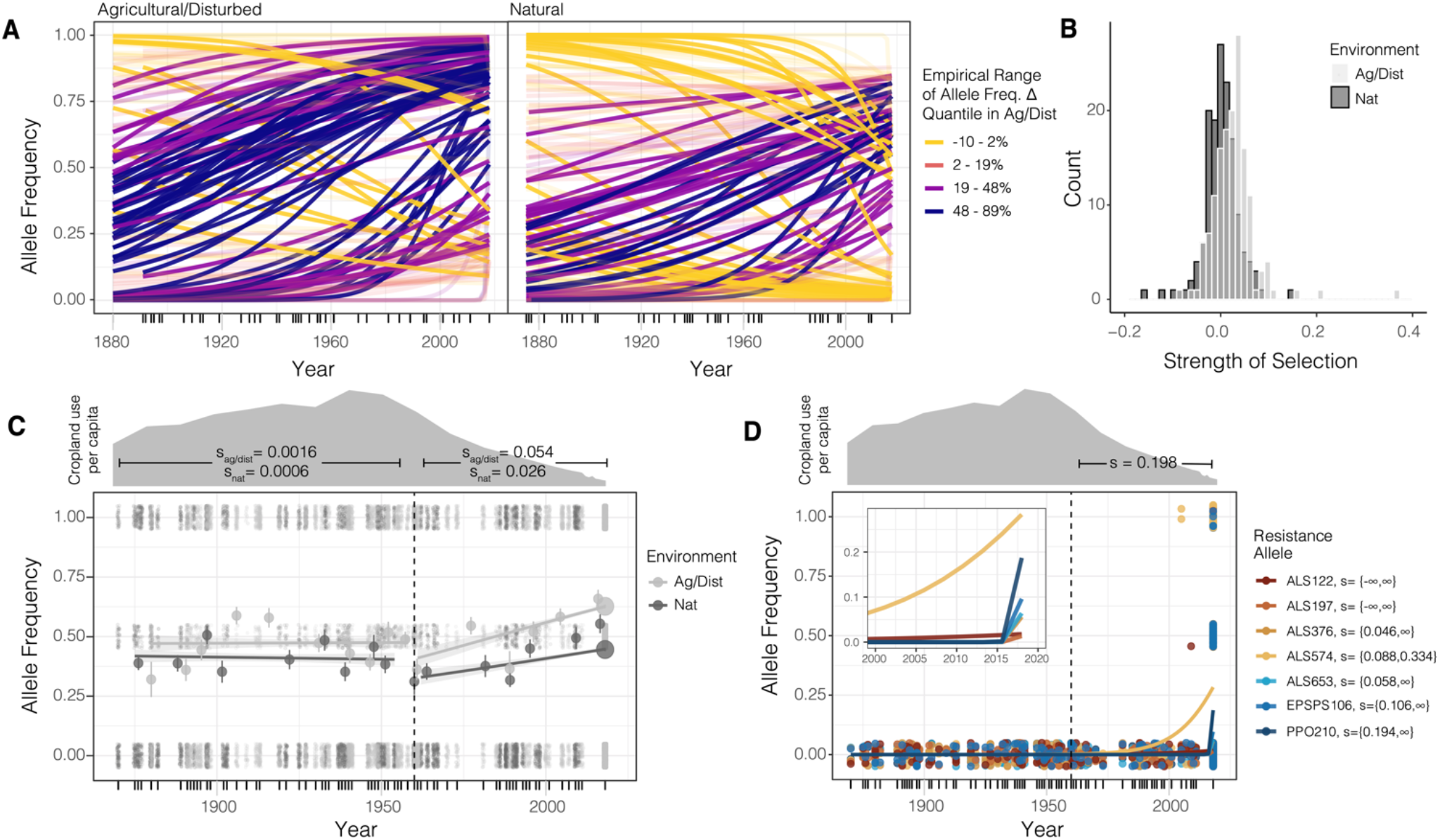
Genomic signatures of agricultural adaptation through time. **A)** Agricultural allele frequency trajectories for each 154 focal SNPs, in agricultural and disturbed habitats (left), and in natural habitats (right). Trajectories colored by the empirical range of the allele frequency change quantile in agricultural and disturbed habitats. Transparent lines indicate those with non-significant evidence of selection at *α*=0.05 after FDR=10% correction. **B)** The distribution of selective strengths on agricultural alleles in natural (dark gray) and agricultural/disturbed (light gray) habitats between 1870 and 2018. **C)** Environment-specific agricultural allele frequency trajectories, before and after the start of agricultural intensification in 1960 (vertical dashed line). Large circles represent moving averages (over both loci and individuals) of allele frequencies, whereas dots represent raw genotype data for each locus and sample from which the allele frequency trajectory is estimated. Cropland use per capita in North America data from (*1*), rescaled by use in 1600, to reflect intensity of agricultural practices. **D)** The trajectory of alleles at known herbicide resistance loci through time, fit by logistic regression for each of the biallelic resistance alleles present in our contemporary data (excluding EPSPSamp with its complex allelic structure). Dots represent genotypes for each historical and contemporary sample at each herbicide resistance locus. 95% confidence interval of the maximum likelihood estimate of selection between 1960-2018 provided in the legend for each resistance allele.

The considerable increase in frequency of these alleles across environments corresponds to remarkably strong selection even when estimated over century-long time periods. The 154 agriculture-associated alleles collectively exhibit a selective strength of 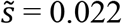 since the 1870s in agricultural and disturbed habitats. However, these alleles exhibit much weaker selection, 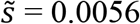, in natural habitats (agricultural and disturbed null interquantile range = [0.0026, 0.0068]; natural null interquantile range = [-0.0018, 0.0018]). An open question in evolutionary biology is what distribution of selection coefficients underlie adaptation (*42*). We estimate that selection on agricultural-associated loci varies between -0.196 and 0.150 in natural habitats, and -0.090 and 0.372 in agricultural and disturbed habitats, reflective of left and right skewed distributions respectively (**Fig 3B, Fig S5)**. The top 15 agriculture-associated alleles that we infer have experienced the strongest selection over the last ∼150 years include SNPs that map near *PPO, ACO1, CCB2, WRKY13, BPL3*, and *ATPD* (**Table S4)**. We find that both the total frequency change of agriculture-associated alleles and the estimated strength of selection in agricultural and disturbed environments are positively correlated with the extent of contemporary linkage disequilibrium around these loci (the number of SNPs with r^2^ > 0.25 within 1Mb) (frequency change: *F = 5*.*16*, p = 0.024, *r = 0*.*12*; strength of selection: *F= 3*.*99*, p = 0.048, *r = 0*.*058*; **Fig S6**), consistent with theoretical expectations for the genomic signatures of recent positive selection (*43, 44*).

We next asked how well the trajectory of modern agricultural alleles reflect the rise of industrialized agricultural regimes across the last century. When we split out samples into those that predate versus those that come after the intensification of agriculture during the Green Revolution, we find that the increase in frequency of agricultural alleles was negligible in agricultural and disturbed environments before the 1960s (predicted 1870-1960 change = 0.005). In contrast, change subsequent to 1960 nearly completely accounts for the observed rise in frequency of modern agricultural alleles (predicted 1960-2018 change = 0.219, versus total 1870-2018 change = 0.221) (**Fig 3C)**. Corresponding estimates of selection by logistic regression using only data from before 1960 shows no evidence of selection on these loci in disturbed and agricultural habitats (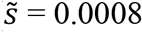, null interquantile range = [-0.0044,0.0020]) or in natural habitats (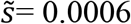, null interquantile range = [-0.004,0.004]). However, samples collected after 1960 reflect a dramatic shift in selection—a collective 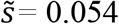 in disturbed and agricultural environments and a collective 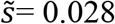 in natural environments (agricultural/disturbed null interquantile range =[0.0064,0.0020]); natural null interquantile range =[-0.0056,0.0054]) (**Fig 3C; Fig S8)**. Together, these results suggest that while most contemporary agricultural alleles were present in historical populations, these alleles only became associated with agricultural and human-managed sites over the last century, on timescales and rates consistent with the rapid uptake and intensification of agrochemicals, controlled irrigation, and mechanization in agriculture.

The historical trajectory of known herbicide resistance alleles epitomizes extreme selection over the last 50 years (**Fig 3D)**. Five out of seven known biallelic herbicide resistance alleles present in our contemporary, paired-environment collections are absent from our historical samples, consistent with the suggested importance of resistance adaptation from *de novo* mutation (*13, 45*) and a particularly recent increase in their frequency. Only three out of 108 historical samples show variation for herbicide resistance, two samples homozygous for resistance at ALS574 and one heterozygous for resistance at ALS122—all of which were sampled after the onset of herbicide applications in the 1960s (**Fig 3D)**. Resolving the very low historical and much higher contemporary frequencies of resistance, we estimate that since the approximate onset of herbicide use in 1960, these seven resistance alleles have collectively experienced a selective strength of 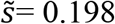 (Z = 2.11, p = 0.035) per year across environment types. Maximum likelihood based estimates of selective strengths for each resistance allele are significant for five of the seven, strongest for PPO210 (s > 0.194), EPSPS106 (s > 0.106), and for ALS574 (s > 0.088) **(Fig 3D; Table S3**).

### Concurrent temporal shifts in ancestry underlie agricultural adaptation

Finally, we explored whether historical demographic change over the last two centuries has played a role in agricultural adaptation. Early taxonomy described two different *A. tuberculatus* varieties as separate species, with few distinguishing characteristics (seed dehiscence and tepal length (*16*)). Sauer’s 1955 revision of the genus, which used herbarium specimens to gauge the distribution and migration of congeners over the last two centuries (*46*), led him to describe an expansion of the southwestern var. *rudis* type (at the time, *A. tamariscinus* (Sauer)) northeastward into the territory of var. *tuberculatus (A. tuberculatus* (Sauer)), sometime between 1856-1905 and 1906-1955. Our sequencing of over 100 herbarium samples dating back to 1828, combined with 349 contemporary sequences (*25, 47*), allowed us to directly observe the change in the distribution of these two ancestral types, adding further temporal resolution to Sauer’s morphological observations of the species’ range shifts, and to assess the role of agriculturally-adaptive standing genetic variation across varieties.

Range-wide, we see clear shifts in the distribution of var. *rudis* ancestry based on fastSTRUCTURE (*48*) inference at K=2 (**Fig S9)** across three-time spans, 1830-1920, 1920-1980, and 1980-2018 (timespan: *F* = 5.47, *p* = 0.0045), and particularly so in the east (timespan x longitude: *F* = 5.49, *p* = 0.0045), consistent with a recent expansion of var. *rudis* ancestry (**Fig 4A)**. Furthermore, we see strong state and province-specific shifts in ancestry through time in our historical sequences (time span by state interaction: *F =* 4.22, *p =* 7 × 10^−5^), highlighting not only the shift of var. *rudis* eastwards (with increases through time in Ontario, Ohio, Illinois, and Missouri) but also the very recent introduction of var. *tuberculatus* ancestry into the most western part of the range in Kansas (**Fig 4B)**. *A. tuberculatus* demography thus appears to have been drastically influenced by human-mediated landscape change over the last two centuries, consistent with the massive recent expansion of effective population size we have previously inferred from contemporary samples over this same timeframe (*45*). That this shift has been most notable over the last 40 years is further consistent with the timescale of agricultural intensification, shifts towards conservation tillage, and rampant herbicide resistance evolution within the species (*19, 45, 49, 50*), suggesting selection on resistance may facilitate the colonization of var. *rudis* ancestry outside its historical range. Along these lines, we find this contemporary range expansion has facilitated the sorting of var. *rudis* ancestry across environments (a longitude by time span by environment interaction: *F* = 5.13, *p =* 4 × 10^−5^; **Fig 4C**), with increasing overrepresentation of var. *rudis* ancestry in agricultural and disturbed environments in the eastern portion of the range through time, as previously suggested (*25*).

**Fig 4.**
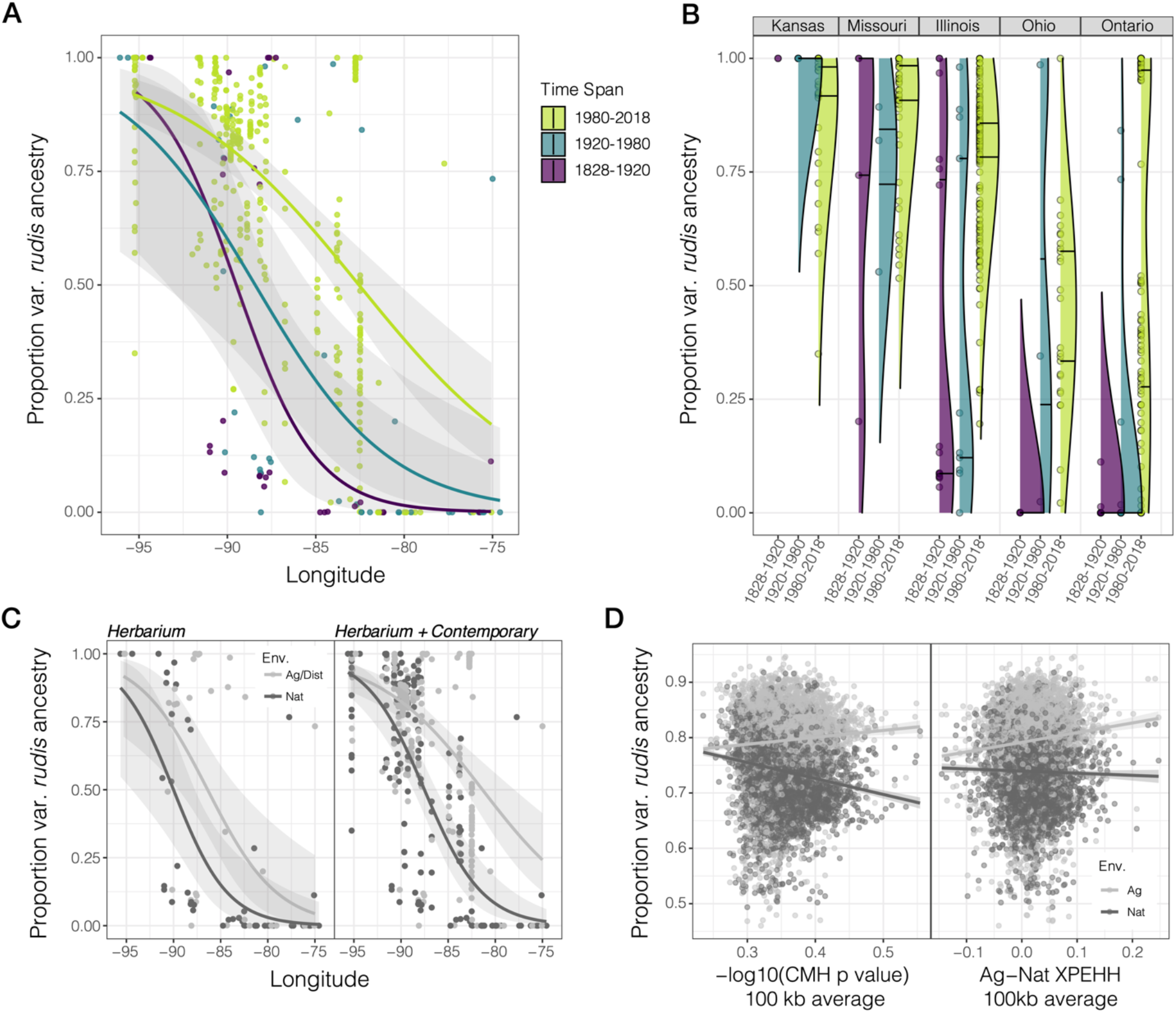
Temporal shifts in the distribution of var. *rudis* ancestry have facilitated polygenic agricultural adaptation. **A)** Longitudinal clines in var. *rudis* ancestry over three timespans, illustrating the expansion of var. *rudis* ancestry eastwards over the last two centuries. In A-C, dots represent individual-level ancestry estimates. **B)** The distribution of individual-level var. *rudis* ancestry by state and through time, illustrating state-specific changes in ancestry. Horizonal lines within each distribution represent first and third quantiles of ancestry. Timespans indicated in (A). **C)** Increasing sorting of individual-level var. *rudis* ancestry into agricultural environments on contemporary timescales. **D)** Environment-specific metrics of selection (CMH p-value and cross-population extended haplotype homozygosity (XPEHH)) across the genome in 100 kb windows positively correlate with var. *rudis* ancestry in agricultural, but not natural habitats (XPEHH by Environment: *F=*9.34, *p=*0.002; CMH by Environment: *F*=99.70, *p* < 10^−16^).

To investigate whether agricultural adaptation has drawn disproportionately from var. *rudis* ancestry, we reconstructed fine-scale ancestry across the genome. Based on analyses in 100 kb windows, we find a least-squares mean of 5.5% (95% CI = [5.0, 5.9%]) more var. *rudis* ancestry genome-wide in agricultural environments compared to the adjacent natural habitat (**Fig S10)**. The environment-specific proportion of var. *rudis* ancestry is not only positively correlated with recombination rate (*F =* 16.67, *p =* 4.5 × 10^−5^, *r =* 0.056) and gene density (*F =* 5.85, *p =* 0.016, *r =* 0.499) but also with SNP and haplotype-based evidence of environment-specific selection. Agricultural, but not natural populations, have an excess of cross-population haplotype homozygosity (agricultural vs. natural XPEHH) and within-pair environmental differentiation (CMH p-value) in genomic regions of high var. *rudis* ancestry (XPEHH by Environment: *F=*9.34, *p=*0.002; CMH by Environment: *F*=99.70, *p* < 10^−16^; **Fig 4D**), implying that ancestry composition genome-wide in large part determines the extent of polygenic agricultural adaptation. These findings suggest that the expansion of var. *rudis* ancestry across the range, particularly in the last 40 years, has facilitated waterhemp’s success in agricultural habitats through providing access to preadapted, standing genetic variation.

## Discussion

Agricultural adaptation in *A. tuberculatus*, a native plant in North America, has occurred over extremely rapid timescales, facilitated by range shifts in response to the agriculturalization of its native habitat. The human-mediated expansion of the southwestern lineage of the species northeastwards since the latter half of the 20th century has introduced new genetic variation across the range, on which selection in agricultural settings could act. Negligible genetic differentiation across habitats in this species refutes the idea of agriculture existing as separate to natural ecosystems (*51*). Despite substantial gene flow, the prevalence of agricultural alleles has increased rapidly since the intensification of agriculture over the last 60 years, in agricultural environments by nearly 6% per year, and even in natural habitats by more than 2% per year. The selective intensity of industrial agriculture is on par with the selective intensities *Arabidopsis* populations in extreme hot and dry environments are predicted to face by 2070 under the high-emissions scenario of climate change (*52*). The effects of agricultural herbicides are even more extreme—range-wide, evolved resistance mutations have on average increased by 20% per year since herbicides were first introduced— permeating even into natural habitats.

While modern, industrial agriculture imposes strengths of selection rarely observed in the wild, this species has in turn escalated the weed management-evolution arms race through a multitude of interdependent mechanisms: range expansion, polygenic adaptation from standing genetic variation, and large effect herbicide resistance mutations. Together, these results highlight that anthropogenic change not only leads to the formation of new habitats but also provides an opportunity for range expansion that may facilitate and feedback with local adaptation, reshaping genetic variation for fitness within native and potentially weedy species.

## Acknowledgements

We appreciate the pivotal contribution of numerous herbaria towards this research, especially the help of Eric Knox at the Indiana University Herbarium, Jamie Lynn Minnaert-Grote at the University of Illinois INHS Herbarium; Tedesse Mesfin at the University of Ohio Herbarium, Anton Reznicek at the University of Michigan Herbarium, Jim Solomon at the Missouri Botanical Gardens, Caleb Morse at the McGregor Herbarium at the University of Kansas, Tyler Smith and Song Wang at Agricultural and Agrifood Canada, and Deb Metsger and Tim Dickinson at the Royal Ontario Museum. We thank Mike Whitlock and Tom Booker (University of British Columbia), as well as Aneil Agrawal and Tyler Kent (University of Toronto), and Ailene Macpherson (Simon Fraser University) for input on the work; Christa Lanz and Rebecca Schwab (Max Planck Institute) for coordinating sequencing of herbarium samples; Ella Reiter (University of Leipzig) for scheduling and coordinating logistics for clean room facility work; and Patricia Lang, Sonja Kersten and Heike Budde (Max Planck Institute) for advice on molecular protocols troubleshooting.

## Author Contributions

JMK, JRS, and SIW conceptualized the paired sampling design, JMK, HAB, DW, JRS, and SIW conceptualized the use of herbarium data, JMK performed contemporary collections and curated the herbarium samples, SML and HAB conceptualized and designed the molecular work with herbarium specimens, SML coordinated the clean room facility work, JMK and SML performed DNA extraction and library preparations of herbarium tissue, SML oversaw the sequencing of herbarium specimens. JMK performed analyses with input from SPO, SIW, and JRS. SPO wrote the migration-selection balance and maximum likelihood models. JMK wrote and revised the paper with inputs from all authors.

## Data and materials availability

All new sequence has been archived at the SRA (BioProject ID PRJNA878842), while scripts and accompanying metadata have been archived on Github at www.github.com/jkreinz/TemporalAdaptation and additionally on Zenodo (*53*).

## Supplementary Materials

This PDF file includes:

Materials & Methods

Figs. S1 to S13

Tables S1 to S4

References 54-68

## Materials and Methods

### Herbarium collections

In 2019, we obtained 10 mg tissue collections of herbarium specimens from 7 herbaria across Canada and the United States and one governmental organization: the Royal Ontario Museum Herbarium, the Museum of Biological Diversity at Ohio State University Herbarium, the Indiana University Herbarium, the Michigan State University Herbarium, the Illinois Natural History Survey Herbarium, Missouri Botanical Gardens, The McGregor Herbarium at the University of Kansas, and Agriculture and Agrifood Canada. We selected samples to have an even representation of habitats through time. Samples were classified as natural (n=54), agricultural (n=28), or disturbed (n=20) based on collectors’ annotations on each plate: any reference to a cultivated field was treated as an ‘agricultural’ collection; descriptions such as dry grassland or riverbank was treated as a ‘natural’ collection; and reference to disturbed soil, railroad tracks, or manicured or managed land was treated as a ‘disturbed’ collection. For inference of contemporary allele frequency and ancestry change through time, samples collected from disturbed habitats were grouped together with the agricultural category—in both of which waterhemp exists as a weed (**Table S5)**. When geographic coordinates were not provided, we referred to the state, county, section, intersection, and landmark descriptions to infer the geographic coordinate of a given sample. In total, we collected samples from 172 specimens, 108 of which were selected for whole-genome sequencing.

### Herbarium DNA extractions & library preparations

The work was performed in the ancient DNA lab at the University of Tübingen. For DNA extraction of the herbarium samples, we followed basic protocol 1 outlined in (*54*). Briefly, under sterile conditions, ∼10 mg of each sample was ground and incubated with N-phenacylthiazolium bromide (PTB)-based mix overnight to lyse DNA. After a shredding step with QIAshredder spin columns, DNA was purified and eluted with DNAeasy Mini spin columns. Sequencing libraries were prepared using the basic protocol 2 outlined in (*54*), performing blunt-end repair, adapter ligation, a fill-in reaction, indexing, and finally PCR amplification (10 cycles) and a cleaning step. The libraries were sequenced on an Illumina NovaSeq instrument on a single flow cell. The sequencing run produced ∼3,442 Gb data, an average of 32 Gb per sample.

### Mapping, damage correction, SNP calling and filtering

We removed adapters, polyQ tails, and merged reads from herbarium sequencing reads using fastp (*55*). Because of the small fragment size of historical DNA, this resulted in a sizable loss of sequence coverage, from 46X coverage to a mean of 11X coverage. Mapping with bwa mem (*56*), we found on average that 89% of merged reads mapped to the female, US Midwestern, reference genome from (*47*), suggesting low rates of contamination by exogenous DNA. Finally, we performed de-duplication of merged reads with DeDup (*57*), which is optimized for merged paired-end sequencing data. This resulted in a final mean per-sample coverage of 9.7X.

We used the program MapDamage (*58*) to quantify damage patterns in the historical DNA. The fraction of C deamination, which leads to C-to-T substitutions, was low, at the first base ∼2% on average across samples, barely inflated above the C-to-T substitution rate across the rest of the reads (**Fig S3**). Nonetheless, the fraction of C-to-T substitutions at the first base was positively correlated with the age of the samples (*r* = 0.46, *t =* -5.31, *p =* 5.94 × 10^−7^; **Fig S3**). We thus used MapDamage to rescale per-base quality scores to take into account the patterns of DNA damage. We called SNPs with freebayes (v1.3.2; with the arguments --use-best-n-alleles 4, --report-monomorphic) in 100 kb regions in parallel across the genome, merged, and then filtered SNPs based on the relationship between QUAL and DP (QUAL/DP > 30). In total, this resulted in 14,139,333 SNPs before merging with our contemporary data and filtering on missing data.

Herbicide resistance alleles in herbarium samples were identified based on known locations of non-synonymous substitutions within ALS, PPO, and EPSPS. Initially, two genotype calls from herbarium samples predated the onset of ALS herbicide use in the 1950s, showing standing variation for resistance at ALS574 and ALS122: one individual heterozygous for Trp-574-Leu collected in 1930 from a sandy agricultural field in St. Louis, Missouri, USA (HB0973); and another individual heterozygous for Ala-122-Ser collected in 1895 from a corn field in Fayette, Ohio, USA (HB0914). Upon further inspection, read-level support for resistance alleles was low with the allelic-bias at these genotype calls being highly skewed (reference to alternate ratio = 1:9 and 2:18, respectively). Similarly, one individual collected in 1967 from the Bottom of Maumee River, Ohio (HB0977) was heterozygous for ALS122, but the alternate resistance allele had support at only one read (reference to alternate ratio =1:7). We subsequently dropped these genotype calls from analyses of selection on herbicide resistance alleles through time. Relatedly, for both the set of 154 focal agricultural-alleles and ancestry informative SNPs used to call fine-scale ancestry across the genome, we investigated the potential for reference bias influencing our estimates of change through time. We calculated mean allelic bias (AB) for each set of SNPs individually for each sample, and asked the extent to which it correlated with collection year (Figure S3). AB for both sets of alleles was not significantly or meaningfully correlated with collection year (AB for agricultural alleles: β =3.987×10^−5^, *p* = 0.316, *t =* 0.7527; AB for ancestry informative alleles: β = -3.877×10^−5^, *p* = -1.882, *t* = 0.0626).

### Metrics of differentiation across Environments: CMH, F_ST_, & XPEHH

To identify fine scale patterns of agrcultural adaptation across the genome, we used the 7,262,599 genome-wide high-quality SNPs called from contemporary agricultural-natural paired populations (n=187 individuals total from 17 pairs of populations, 34 populations in total) from (*25*) (Fig 1). Previously, these data had been only used for genome-wide PCA and faststructure based individual-level ancestry estimates. To make use of our paired sampling design, we used plink (*59*) to perform a Cochran–Mantel–Haenszel test, testing an (environment x SNP | pair) effect after applying a minor allele frequency cutoff of 0.01. We identified candidate agriculturally-adaptive genes based on the nearest gene (bedtools closest) to each LD-clumped, [FDR-corrected] q-value < 0.1 SNP. We found the *Arabidopsis thaliana* orthologues of our *A. tuberculatus* genes with orthofinder (*60*). For genes where orthofinder found no *A. tuberculatus* orthologue and in which our annotation identified no orthologue in closely related species based on gene expression data, we used blastn (*61*) to perform a conclusive search for similar genes across species.

We used plink to calculate Weir and Cockerham’s F_ST_, both between all natural and agricultural samples, and between environments within each population pair, which we later averaged to obtain the mean pairwise F_ST_. For calculation of F_ST_ at the EPSPS amplification, we recoded individuals as 0, 1, 2 based on copy number amplitude (<1.5, 1.5 < copies < 2.5, and >2.5, respectively). Briefly, EPSPS copy number was estimated in Kreiner (2019), through scaling coverage within the EPSPS gene by the genome-wide average. We used selscan (*62*) to calculate the cross-population extended haplotype homozygosity, after read-back and population-level phasing with Shapeit2 (*63*), both of which required knowledge of recombination rates, which we supplied in the format of our imputed LD-based map from (*47*).

### Models of Migration-Selection Balance

Three resistance alleles showed significant differences in allele frequency across natural and agricultural environments (ALS574, EPSPSamp, and PPO210) based on a multiple regression approach (lm: genotype ∼ pair + environment), reflecting differential selection pressures in the face of otherwise high rates of migration (as evidenced by the low genome-wide F_ST_). Indeed, previous experimental work on costs on resistance has shown several of these herbicide resistance mutations to be associated with substantial costs: the ALS574 mutation has been associated with a 67% reduction in above-ground biomass in *A. palmeri* (*64*), whereas the EPSPS amplification has been associated with a 25% reduction in dry biomass in *A. tuberculatus* (*39*) but is associated with no observed cost in *A. palmeri* (*65*). In the context of the experimental conditions and genotypes used in Wu et al., 2019 (*66*), no costs of the PPO210 deletion were found. However, not accounting for realistic genotypic and environmental heterogeneity is a major limitation of experimental approaches to assaying costs (*40*).

We take a model-fitting approach that implicitly takes into account such heterogeneity by looking over a diverse set of genotypes, environments, and selective regimes. Specifically, we fit a two-patch, two-allele model of migration-selection balance to estimate the relative magnitude of migration and selection across environment types. In each patch, we first assume that the life cycle starts at the adult stage, followed by migration and then selection among the juveniles to the next census among adults:

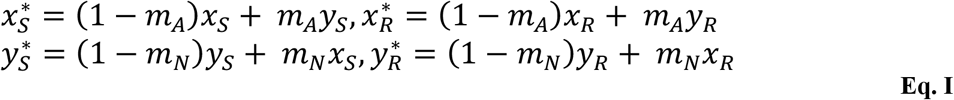

where *m*_*N*_ and *m*_*A*_ represent immigration rates of alleles into natural and agricultural sites, respectively; 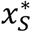 and 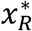 represent the frequency of the susceptible and resistant allele in agricultural environments after migration; and 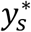 and 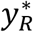 represent the frequency of the susceptible and resistant allele in natural environments after migration. Assuming random mating, the frequencies of resistant (in agricultural, 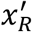; in natural, 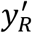) and susceptible alleles (in agricultural, 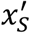; in natural, 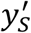) after selection are proportional to:

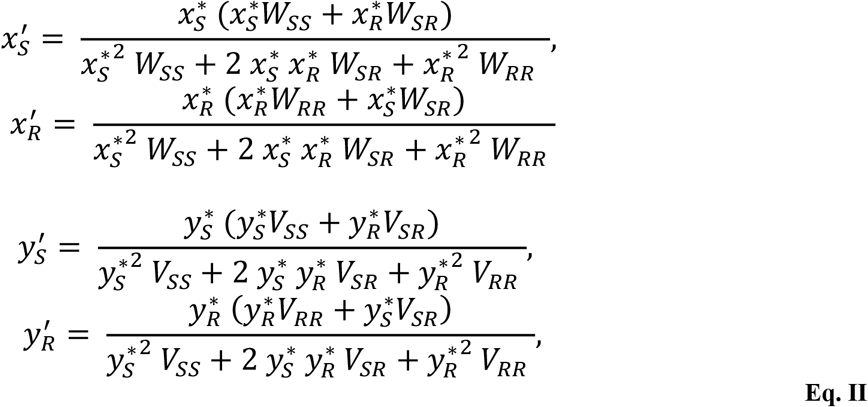

where *W* and *V* reflect the average fitness of each genotype in agricultural and natural environments, respectively. Assuming additivity with *s*_*N*_ measuring the selective cost of the resistant allele in natural environments (*V*_*RR*_ = 1 − *s*_*N*_, *V*_*RS*_ = 1 − *s*_*N*_/2, *V*_*SS*_ = 1) and *s*_*A*_ measuring the selective benefit of the resistant allele in agricultural environments (*W*_*RR*_ = 1 + *s*_*A*_, *W*_*RS*_ = 1 + *s*_*A*_/2, *W*_*SS*_ = 1) and assuming and that migration at the loci is weak (*m <<* 1), a given pair of populations is expected to approach a steady state, where:

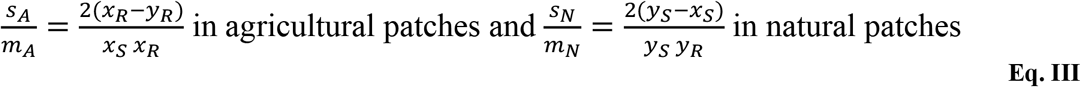

While it is not possible to solve for selection directly in the absence of data on migration rates, these formulae allow us to estimate the strength of divergence by inferring the strength of selection relative to migration in natural 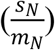 and agricultural 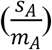 environments, as presented in **Table S3**. The ratio of these metrics gives the ratio of the cost faced per migrant that has arrived in natural environments versus the benefit per migrant that has arrived in agricultural environments, assuming that the pair of populations is near equilibrium. It is worth emphasizing the “per migrant” nature of these measures (*s*_*N*_/*m*_*N*_ versus *s*_*A*_/*m*_*A*_); even if the selective benefit of resistance in agricultural environments were stronger than its cost in natural environments (*s*_*A*_ ≫ *s*_*N*_), it is possible for the benefit of resistance per migrant to be less than its costs per migrant (*s*_*A*_/*m*_*A*_ ≪ *s*_*N*_/*m*_*N*_) when migration rates are higher into agricultural environments (*m*_*A*_ ≫ *m*_*N*_). We also note that the approach to migration-selection balance occurs exponentially at a rate proportional to the selection coefficient (when *m*_*A*_, *m*_*N*_ ≪ *s*_*A*_, *s*_*N*_ ≪ 1) and so should occur rapidly at sites under strong selection (**Supplemental Index 1**).

### Logistic models of temporal allele frequency change

We used CMH outliers from the contemporary paired population scan to investigate patterns of agricultural-allele frequency change over the last two centuries. We were interested in tracking independent allele frequency trajectories, so from the 403 SNPs with CMH p-values that exceeded 10% FDR correction (*p* < 6 × 10^−6^), we performed a subsequent clumping step, effectively retaining a set of largely unlinked SNPs (**Fig S11)** that represent the most significant SNP in a particular region of the genome. Specifically, we used plink --clump, to find the most significant hit genome-wide, scan 1 Mb around it, and exclude any SNP from the resulting output that is associated with the focal SNP with r^2^ > 0.25. This algorithm was repeated until all SNPs passing the genome-wide significance threshold had been clumped. This resulted in 251 loci that on average showed a 17.9% allele frequency difference between extant agricultural and natural environments. Average LD across focal SNPs was 0.043, with only four pairs of SNPs showing high pairwise LD (r^2^ > 0.4) with another SNP. All of these four SNP pairs are found on separate chromosomes from the SNP with which it has high LD, suggesting the correlation is driven by selection and migration not linkage (alternatively, genome-assembly or polygenic adaptation [such as in (*45, 67*)] may drive such a signal). At each SNP, we labelled the alleles such that the agriculture-associated allele was the one whose frequency was higher in agricultural sites than natural sites.

We then found the intersection of these modern agriculture-associated alleles, identified in our contemporary paired collections, with the historical, filtered SNPs from the herbarium sequence data. 154 loci were present in the historical samples with the same reference/alternate allele combinations. Because the definition of agriculture-associated alleles depends on their relative frequency across environment types, such alleles include both reference (91/154) and alternate (63/154) bases. We extracted a matrix of 0, 0.5, and 1 values, representing the frequency of the agricultural allele for each locus within each individual, for samples from both our contemporary and historical collections. Combining these individual agricultural allele frequencies at each locus across historical and contemporary datasets, we then performed a logistic regression in R (glm function, family=“binomial”) of genotype on collection year, separately on samples from either natural or agricultural and disturbed environments. From each logistic regression, we extracted the logit-transformed slope, p-value, and standard error, as well as the predicted value (allele frequency) for the years 1870 and 2018, representing the minimum sample year and maximum sample year. While we have samples dating back to 1828, we constrained this analysis to samples collected after 1870, as the density of samples before then is low (n=4), with no representation of samples from agricultural environments. The slope of the logistic regression gives an estimate of the selection coefficient, *s* (specifically, slope = *s*/2) where *s* is the difference in fitness between the two homozygotes and the division by two comes from using a diploid model. This selection coefficient estimates the net fitness benefit (if *s*>0) of the agriculture-associated allele, as determined by the rise in frequency over time of that allele. Specifically, with homozygote fitness of 1+*s* and heterozygote fitness of sqrt(1+*s*)∼1+*s*/2, measured relative to the wildtype homozygote, the allele frequency over time has a generalized logistic form:

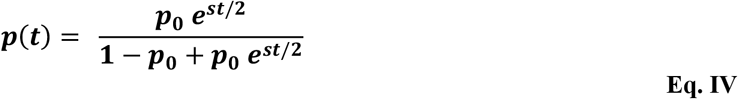

where *p*_0_ is the allele frequency at time *t* = 0.

The total allele frequency change at each locus was calculated by taking the difference between the predicted frequency of the allele in 2018 and 1870. We merged the output of these locus-specific logistic regressions in agricultural environments, with both SNP and haplotype-based statistics from these same individuals to identify contemporary correlates of the magnitude of allele frequency change and selection through time. Specifically, we examined how well contemporary recombination rate, XPEHH, CMH p-value, number of SNPs in linkage disequilibrium (r^2^ > 0.25) with the focal SNP (< 1Mb; i.e., number of SNPs in a clump), and distance between linked SNPs, explained both the total allele frequency change and the estimated strength of selection (**Fig S6)**.

We also performed a separate set of analyses, where a logistic regression was used to analyze the trajectory of all agricultural alleles or known herbicide resistance alleles at once, first across samples from natural environments and then for samples from agricultural and disturbed environments (‘genotype ∼ year + locus’). We further partitioned samples in each environment to those that predate or are subsequent to the 1960s, to infer the importance of the intensification of agriculture and herbicides in shaping the strength of selection on contemporary agricultural alleles (**Fig 3C, D)**. For each of the four logistic regressions ran on these partitioned sets of data, the slope of the year term represents a joint estimate of the strength of selection for agricultural alleles, between 1870-1960 or 1960-2018, in natural or agricultural environments. We refer to this joint estimate of selection at multiple loci as 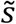.

To test whether a shift in selection at 1960 was statistically supported, we also compared our full model analyzing temporal signatures of allele frequency change between 1870-1960 to one that fits either two or three logistic regression lines across that time frame (i.e., a segmented logistic regression). A segmented logistic regression with two breakpoints provides the best fit to our data, compared to a model with either one or no breakpoints (two-break segmented AIC=54360.55, one-break segmented AIC =54437.66, non-segmented AIC=54444.67), and converges on 1913 and 1961 breakpoints, the later supporting a priori hypotheses and our interest in interrogating signals before and after the start of the Green Revolution in 1960 (**Fig 3C)**. For comparison and visualization purposes of shifts in the trajectory of selection across these time periods, we also present the splines of environment-specific cubic model fits in **Fig S7**.

We designed a randomization test to model the expected distribution of allele frequency change through time under null processes (drift, migration, selection, and demographic change). In particular, we were interested in quantifying the potential bias associated with high frequency alleles having more leeway to change more through time, as compared to a set of lower frequency alleles. We thus randomly sampled 154 loci across the genome from our contemporary collections (the same number as our observed clumped and historically matched set of agricultural alleles), repeating this sampling 1000 times. Importantly, each sample exactly matched the frequency distribution observed for extant agricultural alleles (**Fig S4**). This randomization was done independently in each environment, such that the alleles sampled to match the extant agricultural-allele frequency distribution in agricultural environments in one iteration were different from the alleles sampled to match the frequency distribution in natural environments. To account for the ascertainment bias in our set of putatively agriculturally adaptive alleles—alleles that show the greatest excess of allele frequency in agricultural compared to natural environments—we similarly defined alleles at each randomly drawn SNP (i.e., the agriculture-associated allele was the one found at greater frequency in agricultural sites than in natural sites, whether it was the reference or alternate allele). In each of the 1000 randomizations within each environment, we then performed the same analyses as above: matching these alleles in our historical samples, producing a matrix of genotype data for both contemporary and historical sets, and performing a logistic regression for each locus, as well as logistic regression on all loci at once, for either samples from natural or agricultural environments, and for those that either preceded or were subsequent to 1960. Using the 1000x randomizations, we then computed the 2.75 and 97.25% quantiles (“null 95% interquantile range”) of the statistics of interest (total allele frequency change and selection coefficients) to compare against our observed values. Note that these null expectations implicitly account for changing ancestry through time, as genome-wide genotypes reflect the spread of var. *rudis* over time and space.

We performed forward-time simulations in SLIM (v3.7.1) to validate the robustness of our space-time herbarium sampling approach for inference of the strength of selection for a set of alleles. On a genomic background of length 100kb, with a recombination rate scaled to approximately that of *Arabidopsis thaliana* (4×10^−6^), we started by evolving additive mutations (with a mutation rate of 5×10^−6^) neutrally for 2000 generations in 5000 individuals. After this time period, we imposed an environmental shift causing those previously neutral mutations to become beneficial [with an exponential distribution of fitness effects centered on *s*=0.1]. This scenario represents selection on standing genetic variation and, potentially, any new mutations that arise after the onset of selection. After this environmental shift, we start our temporal sampling following the same temporal distribution and total sample size used in our manuscript (i.e., *t* = 0, 5, 6, 7, 10, 10, 10, 12 … 141, representing years sampled after 1870, *n* = 104 [removing four individuals from < 1870]), with individuals randomly sampled across 2D space. From these simulations, we find that our temporal and spatial historical sampling approach is able to provide an accurate estimate of *s*. On average across 500 simulations, the correlation coefficient between estimated *s* and true *s* was 0.61 (SE=0.067). Thus, the method employed to estimate the strength of selection is expected to be accurate for the sample sizes in this study, although we expect accuracy to decline with smaller sample sizes, weaker selection, and/or smaller species-wide effective population sizes.

### Maximum likelihood estimate of selection

For known biallelic herbicide resistance alleles (excluding only the complex EPSPS amplification), we were particularly interested in understanding selection on each allele over time. We used a maximum likelihood approach to estimate the strength of selection for each resistance allele between 1960-2018, along with a 95% confidence interval using profile likelihood. Summing overall years (*t*), the log-likelihood of observing the data is given by the binomial sampling formula describing the chance of observing the number of resistant (*n*_*R*_) and susceptible alleles (*n*_*S*_) in any given year:

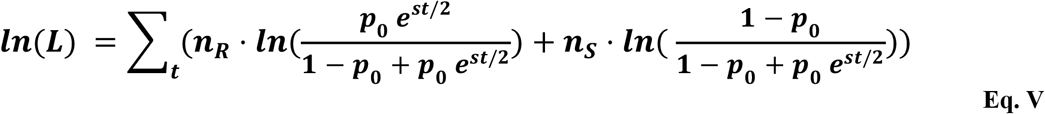

where *p*_*0*_ represents the frequency of the allele when *t* = 0 (defined as the present for ease of computation) and *s* represents the strength of selection (see logistic allele frequency trajectory equation above), both of which are unknown and estimated by maximizing the likelihood. Because many of the resistant alleles were only observed in contemporary samples, selection must be sufficiently strong on recent timescales to explain this rise, but any larger value of selection is also able to explain the data (i.e., the likelihood surface becomes flat with increasing *s* in several cases). Because maximum likelihood points are poorly estimated when the likelihood surface becomes flat, we only present the 95% confidence interval in the main text (i.e., those values of *s* for which the ln(L) falls within 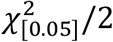 of the maximum likelihood), but present maximum likelihood estimates in Table S3 as well. We implemented this algorithm in R, using the mle2 function implemented within the bblme package in R.

### Ancestry inference

For genome-wide ancestry inference, we merged filtered SNPs from herbarium samples with high-quality SNP sets from (*25*) (n=187, collections from 2018) and (*47*) (n=162, collections from 2015), resulting in 457 individuals and representing all resequenced *A. tuberculatus* whole genomes (*n* of SNPs = 1,269,007). We used faststructure (*48*) to infer individual-level ancestry, taking the proportion of an individual’s assignment to a grouping at K=2 to represent either var. *rudis* or var. *tuberculatus* ancestry. An individual’s proportion of var. *rudis* ancestry was then analyzed in a multivariate regression that tested how well var. *rudis* ancestry was explained by longitude, latitude, environment (natural or agricultural), timespan (1800-1920 [n=39], 1920-1980 [n=44], 1920-2020 [n=374]), a two-way timespan by longitude interaction, a two-way timespan by state interaction, and a three-way timespan by environment by longitude interaction (Individual ancestry assignment ∼ longitude + latitude + environment + timespan + timespan:longitude + timespan:state + timespan:environment:longitude)

We also used plink to perform a principal-component analysis of merged SNPs from just herbarium samples (**Fig S12)** and all 457 samples jointly (**Fig S13)**.

We were interested in the distribution of var. *rudis* ancestry across the genome, and so used LAMP (*68*) to assign ancestry to SNPs, based on two reference populations homogenous for either var. *rudis* or var. *tuberculatus* ancestry (Kansas and Ontario Natural Populations, respectively; (*47*)). Ancestry informative SNPs were those with an F_ST_ > 0.40 (2x the mean genome-wide ancestry differentiation between varieties, in these two populations) between these reference populations and that were also in common between datasets (<20% of samples with missing data) after merging historical sequences with the contemporary paired sequence data (*25*). Since LAMP requires recombination rate information, we also imputed the LD-based genetic map from (*47*) to the ancestry-informative SNPs to get genetic distance between each. Finally, we performed the LAMP analysis, one population at a time, one scaffold at a time. After merging SNP-wise ancestry assignments across scaffolds, we calculated the mean, 5%, and 95% quantile of var. *rudis* ancestry in 100 kb regions for each population, and eventually, each environment (**Fig S10)**.

To understand the relationship between ancestry, agricultural selection, and genomic architecture, we performed a multiple regression to quantify drivers of fine-scale ancestry across the genome. We regressed the individual proportion of var. *rudis* ancestry in 100 kb windows across the genome against gene density, recombination rate, scaffold, environment, average CMH score, average XPEHH (difference in extended haplotype homozygosity across environments), the interaction between environment and average CMH score in each window, and the interaction between environment and the mean XPEHH in each window (100kb mean ancestry ∼ scaffold + mean gene density + mean recomb + mean xpehh:env + mean cmh:env + env). The least squares effect of environment on ancestry was taken to calculate the average difference in ancestry between agricultural and natural environments.

**Fig S1.**
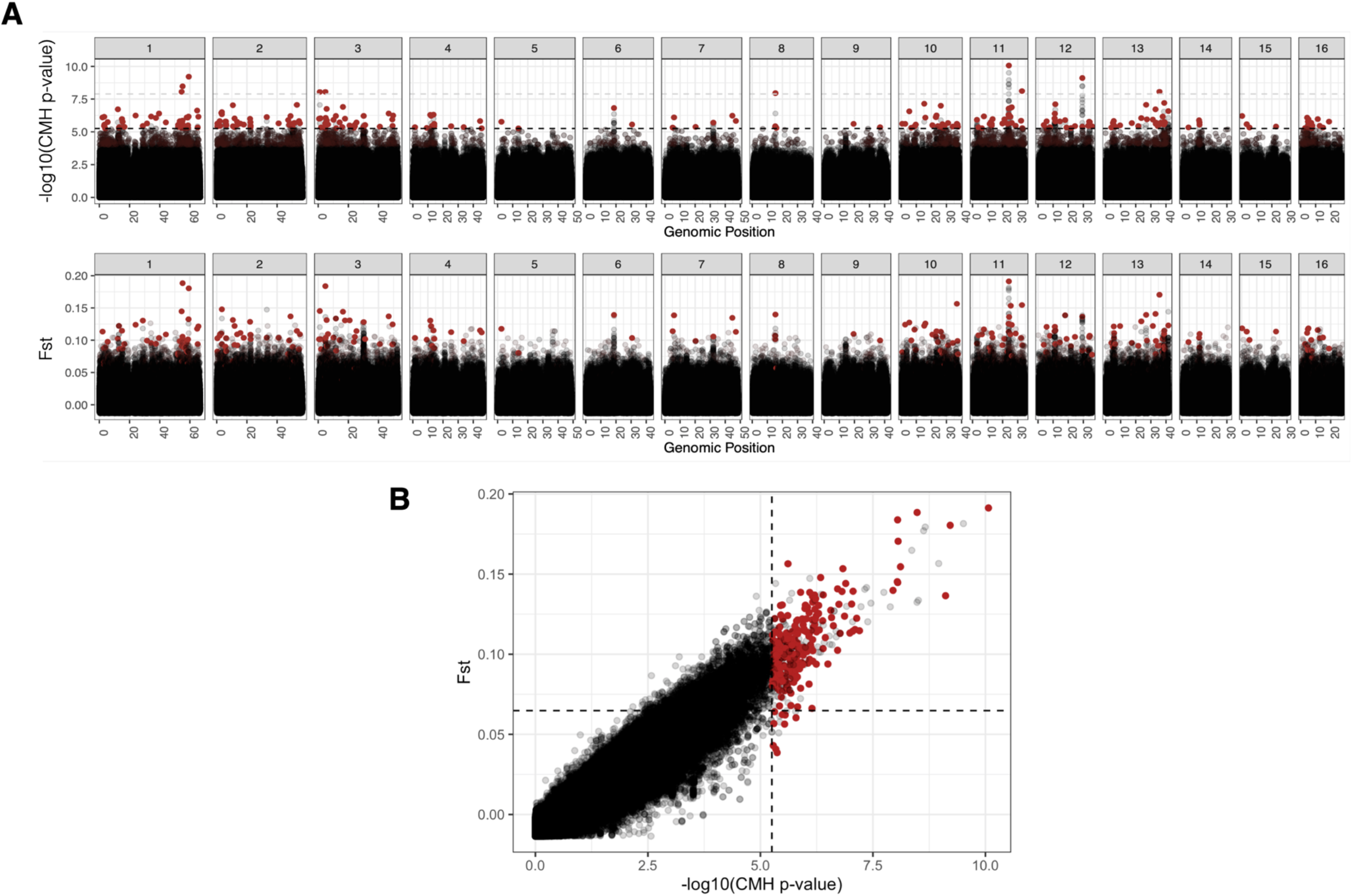
Strong congruency between results of a CMH genome-wide scan (assessing environmental differences stratifying for population pair) versus a between-environment F_ST_ genome-wide scan (differentiation among individuals pooled within natural environments and within agricultural environments). A) Two Manhattan plots showing the distribution of CMH -log_10_(p-values) [top] and Fst values [bottom] at SNPs across the genome. B) Between-environment F_ST_ is plotted against the CMH -log_10_(p-values), showing a strong correlation (Spearman’s *ρ* = 0.905; Pearson’s *r* between Fst and CMH χ^2^ = 0.957). In both A and B, red dots indicate clumped, putative agriculturally adaptive SNPs as inferred from the CMH scan after applying a 10% FDR cutoff.

**Fig S2.**
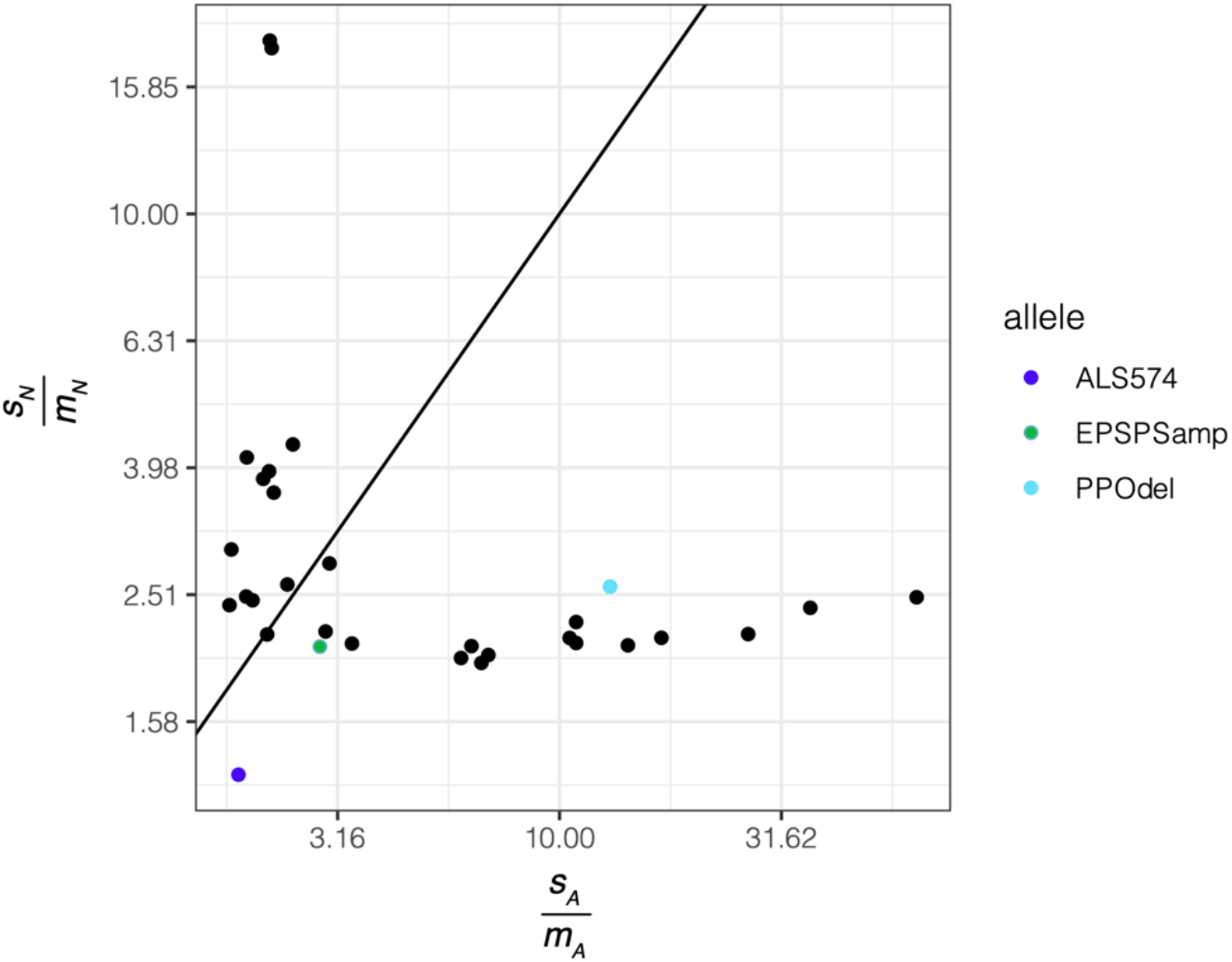
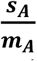 (representing selective benefit per migrant in agricultural habitats) versus 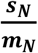 (representing the selective cost per migrant in natural habitats) for the 30 independent loci with the most significant CMH scan hits, compared to the 3 common herbicide resistance alleles with significantly different allele frequencies among natural and agricultural environments. Diagonal line represents equal agricultural benefits compared to natural costs, scaled by migration.

**Fig S3.**
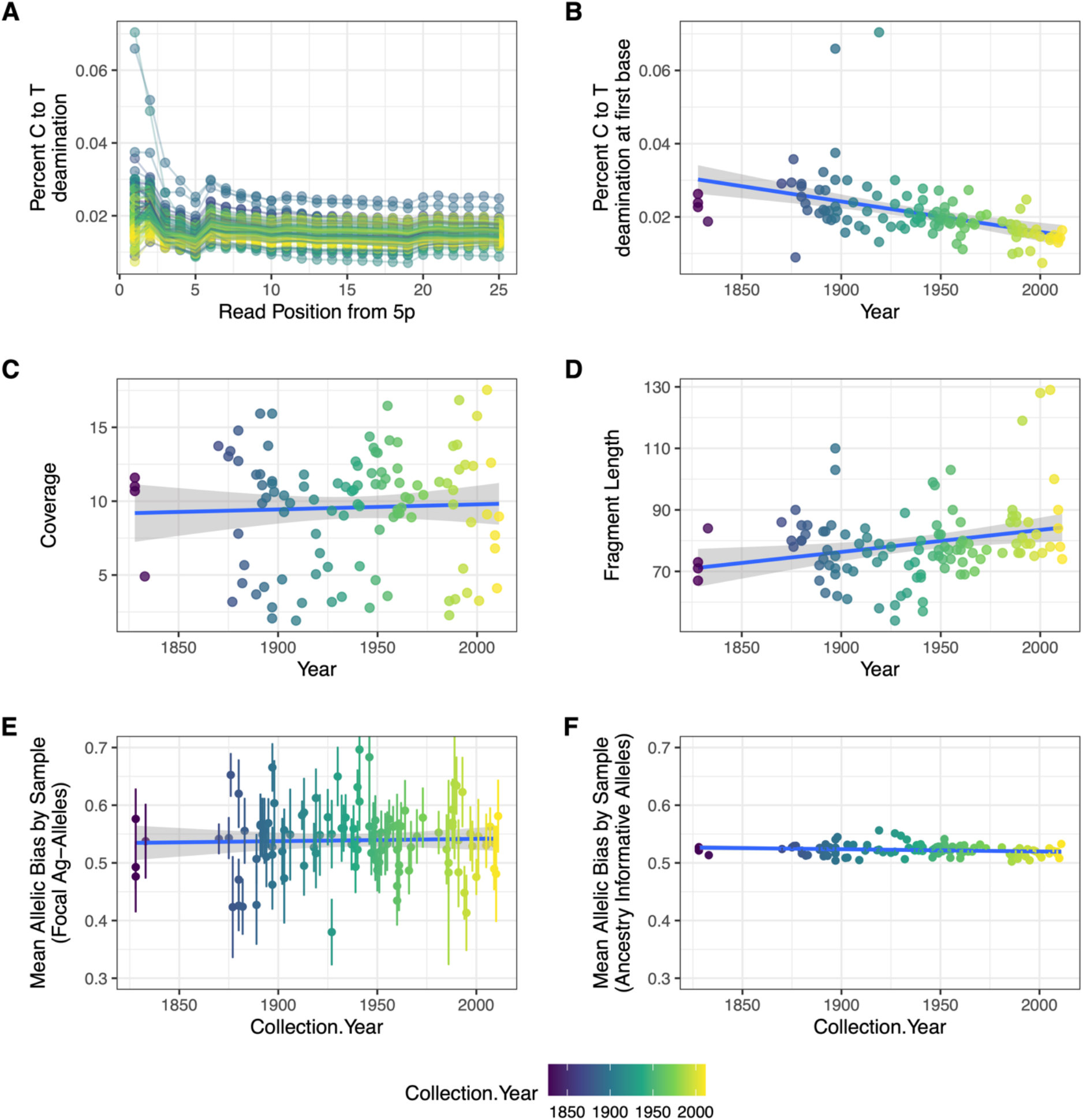
Percent C-to-T deamination by read position (A), along with the correlation of collection year with percent C-to-T deamination at first base (B). C-D represent temporal correlates with genome-wide coverage (C), fragment length (D), mean allelic bias by sample for focal agriculture-associated alleles (E) and mean allelic bias by sample for ancestry informative alleles (F). For B-F, each dot represents sample-wise means for each108 sequenced herbarium specimens.

**Fig S4.**
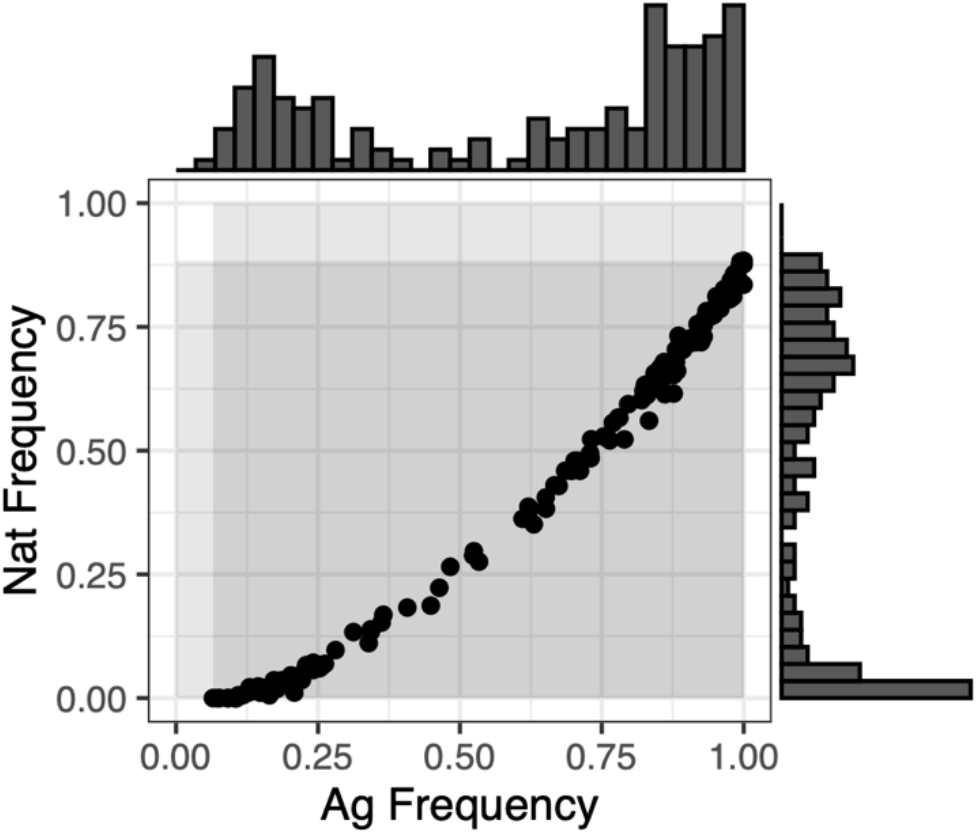
The distribution of frequencies for agricultural-associated alleles in agricultural samples along the x-axis, and in natural samples along the y-axis. Null distributions for an expectation of change in the frequency in our focal set of alleles was generated by producing randomized allele sets of the same size (n=154) matching the extant frequency distributions shown here, first in natural environments (top histogram), and then in agricultural environments (right histogram).

**Fig S5.**
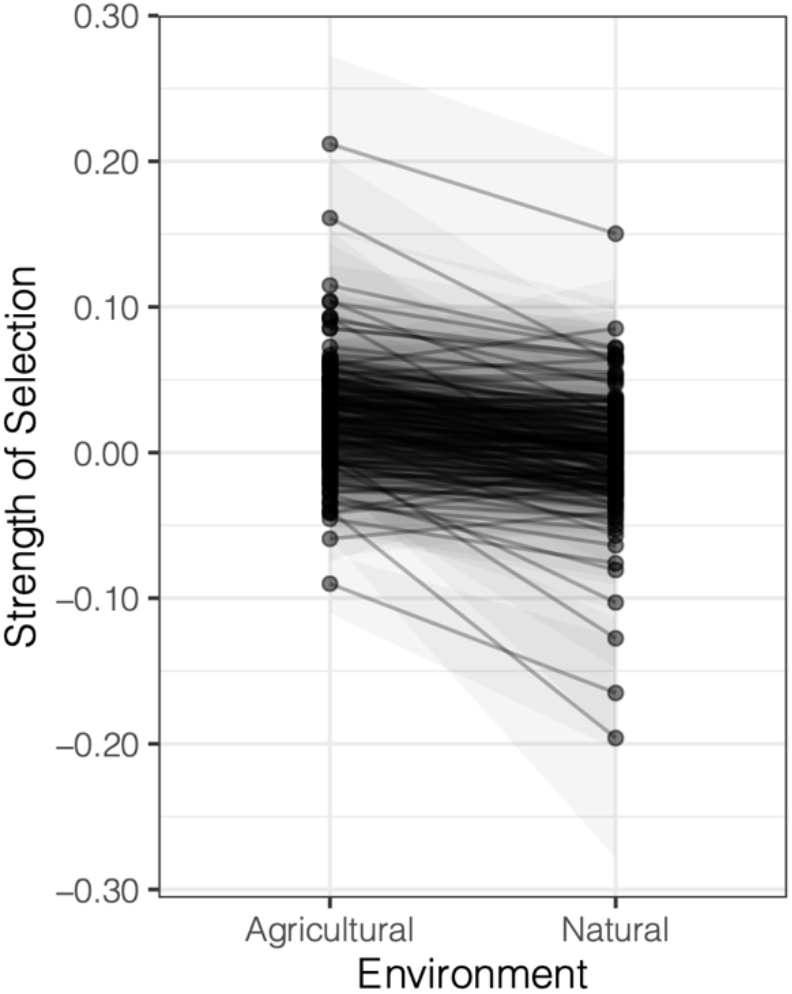
Inferred strength of selection on 154 agricultural alleles through time, in either agricultural or natural environments. Selection coefficients were extracted from logit-transformed logistic regressions of genotype on year, run separately for each locus in each environment. Gray ribbon for each locus represents the bounds of the standard error associated with the estimate of selection in each environment.

**Fig S6.**
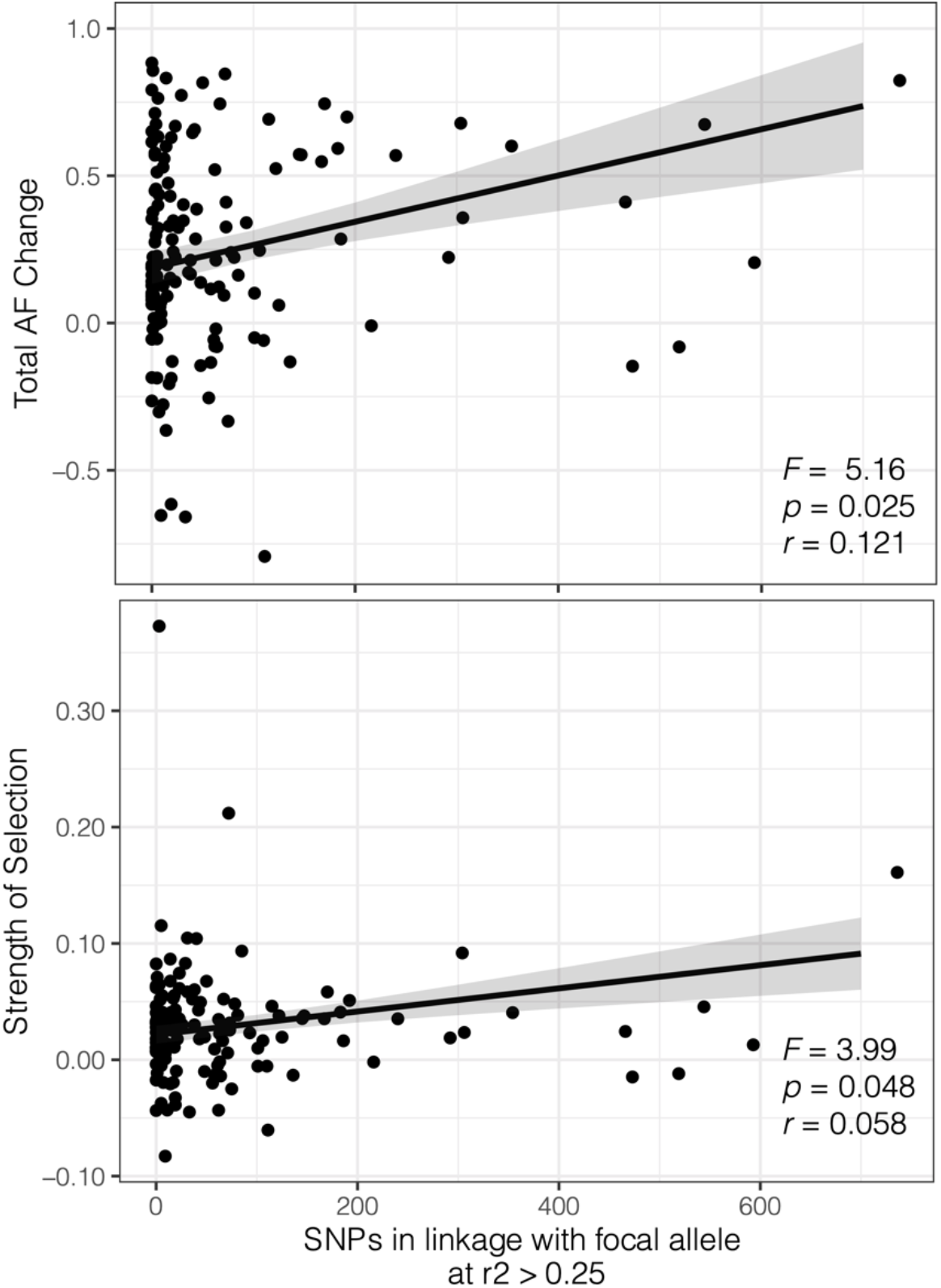
The association between contemporary patterns of linkage disequilibrium (number of SNPs within 1Mb of focal agricultural-associated allele with r^2^>0.25) and allele frequency change (top) or selection (bottom) observed over the last 150 years across herbarium samples. Regression line shows the least square mean effect of contemporary associations from a multiple regression analysis.

**Fig S7.**
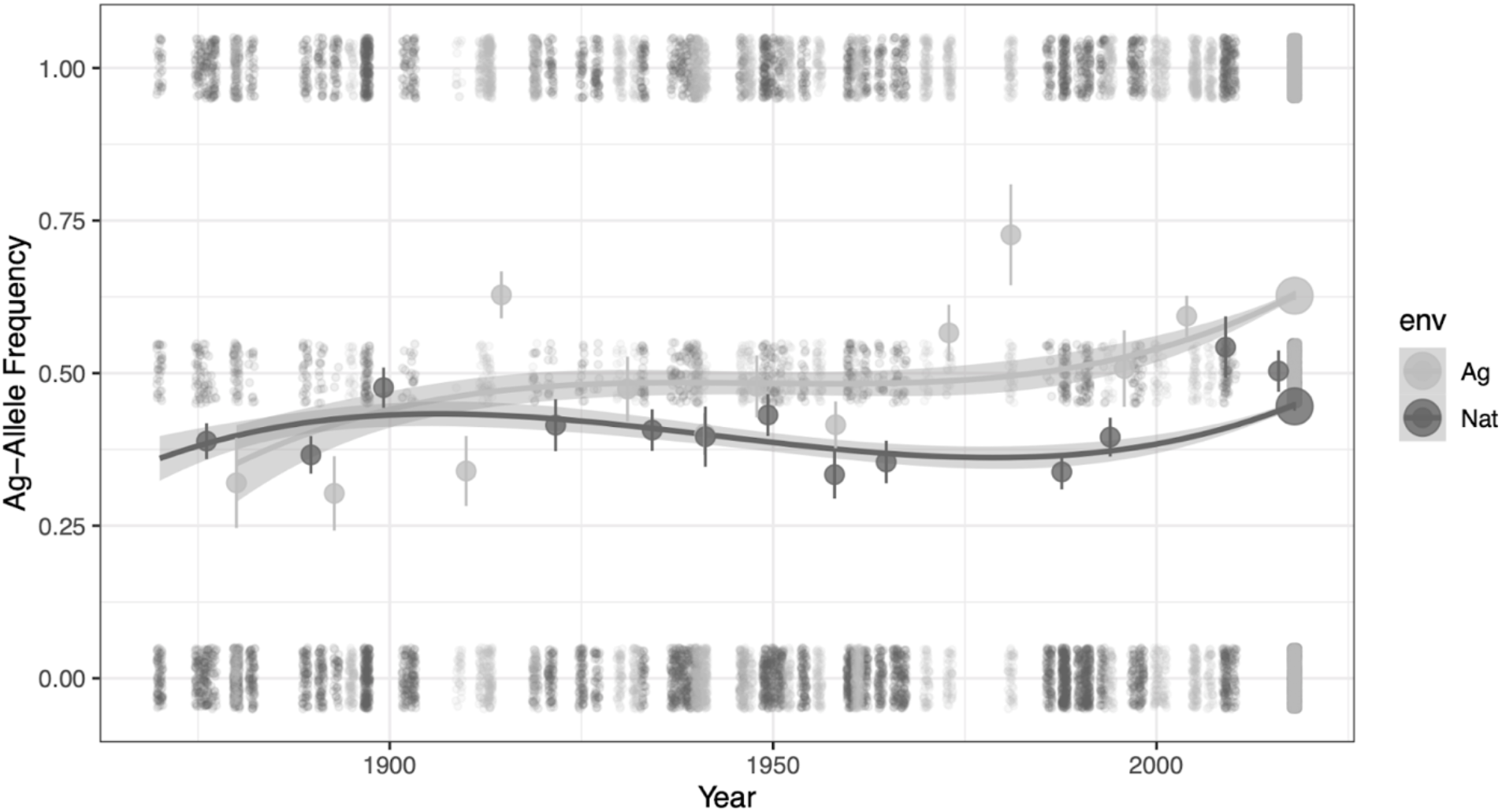
Cubic splines that illustrate the environment-specific frequency change of modern agriculture-associated alleles through time since 1870. Gray ribbon denotes the 95% CI.

**Fig S8.**
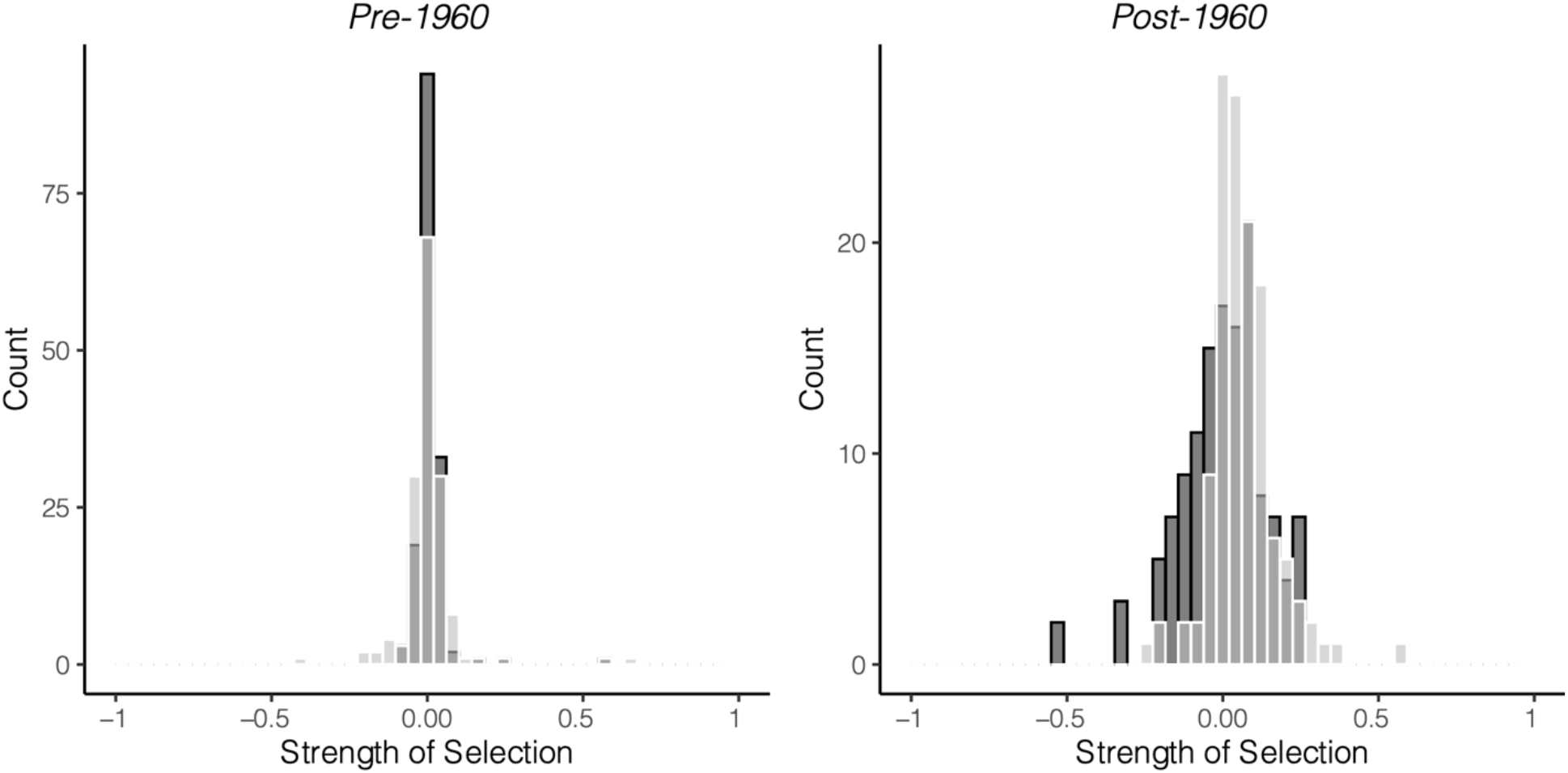
Overlayed histograms of logistic-model estimates of selection before (left) and after (right) the 1960s, the start of agricultural intensification, for agriculturally-associated alleles in natural (dark gray) versus agricultural and disturbed (light gray) environments (medium gray occurs when counts overlap).

**Fig S9.**
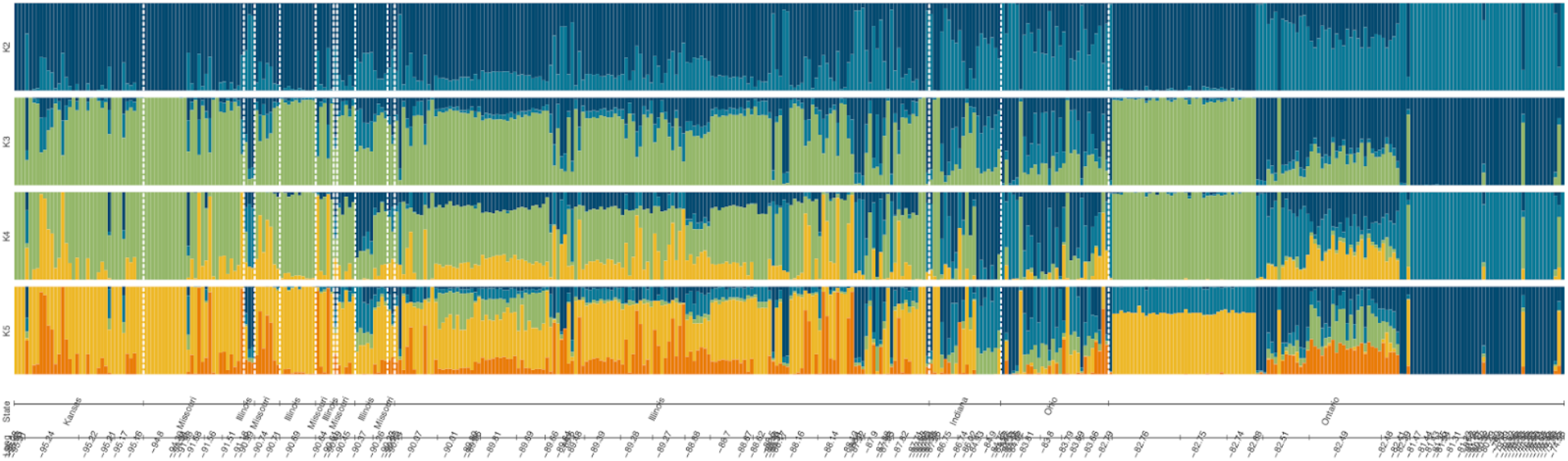
Longitudinal and state-wise patterns of ancestry across 457 *A. tuberculatus* individuals from contemporary and historical sampling, inferred from faststructure. Samples sorted by longitude, from west (left) to east (right). White dashed lines denote clusters of specimens sampled from different states and provinces across this longitudinal gradient. K=2 taken as var. *rudis* versus var. *tuberculatus* ancestry, as in (*47*).

**Fig S10.**
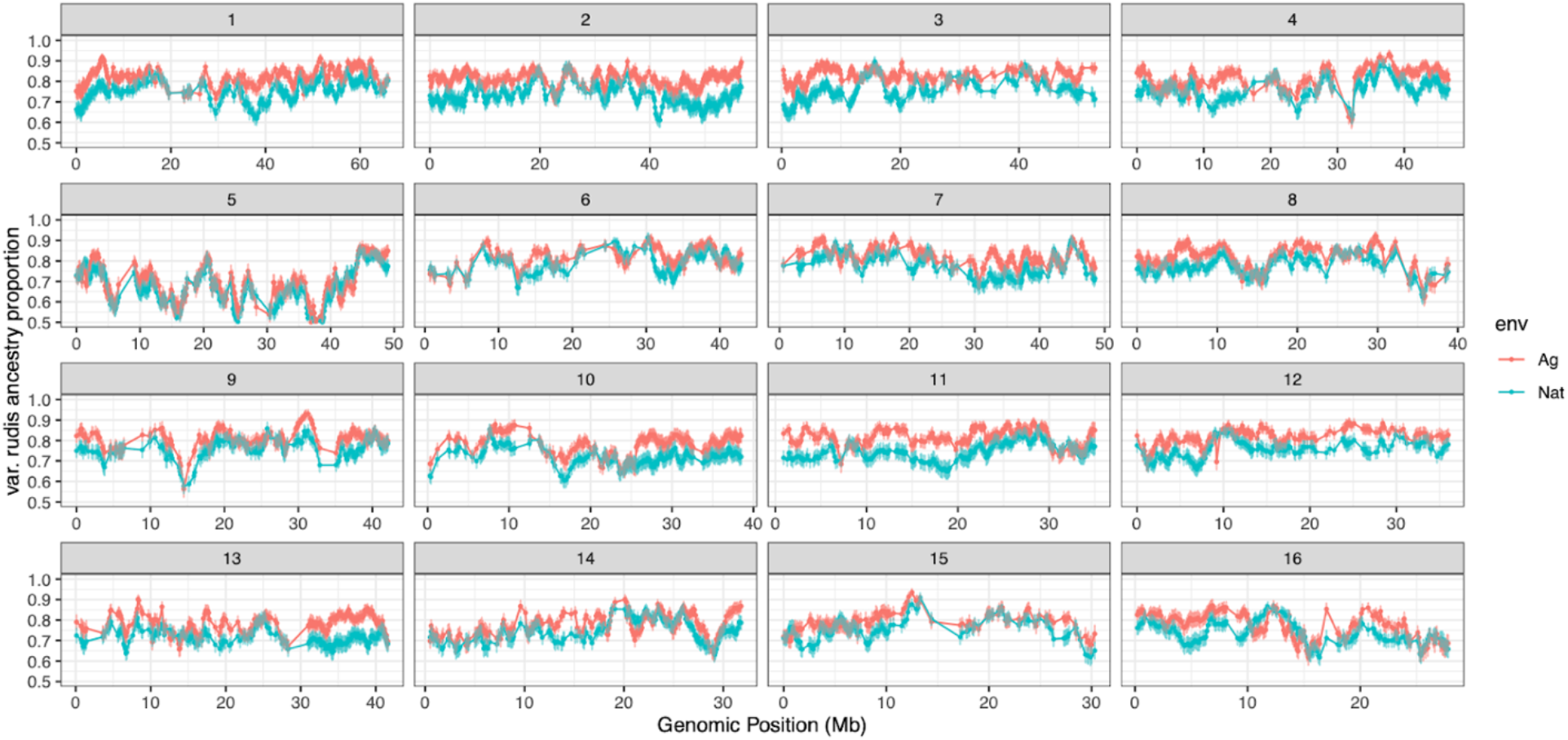
Excess of var. *rudis* ancestry in agricultural compared to natural environments, in 100 kb regions across the genome. Lines depict the mean ancestry across all populations within each environment, with error bars showing the mean 5th and 95th percentile of ancestry across populations. Fine-scale ancestry estimates were inferred with LAMP (*68*).

**Fig S11.**
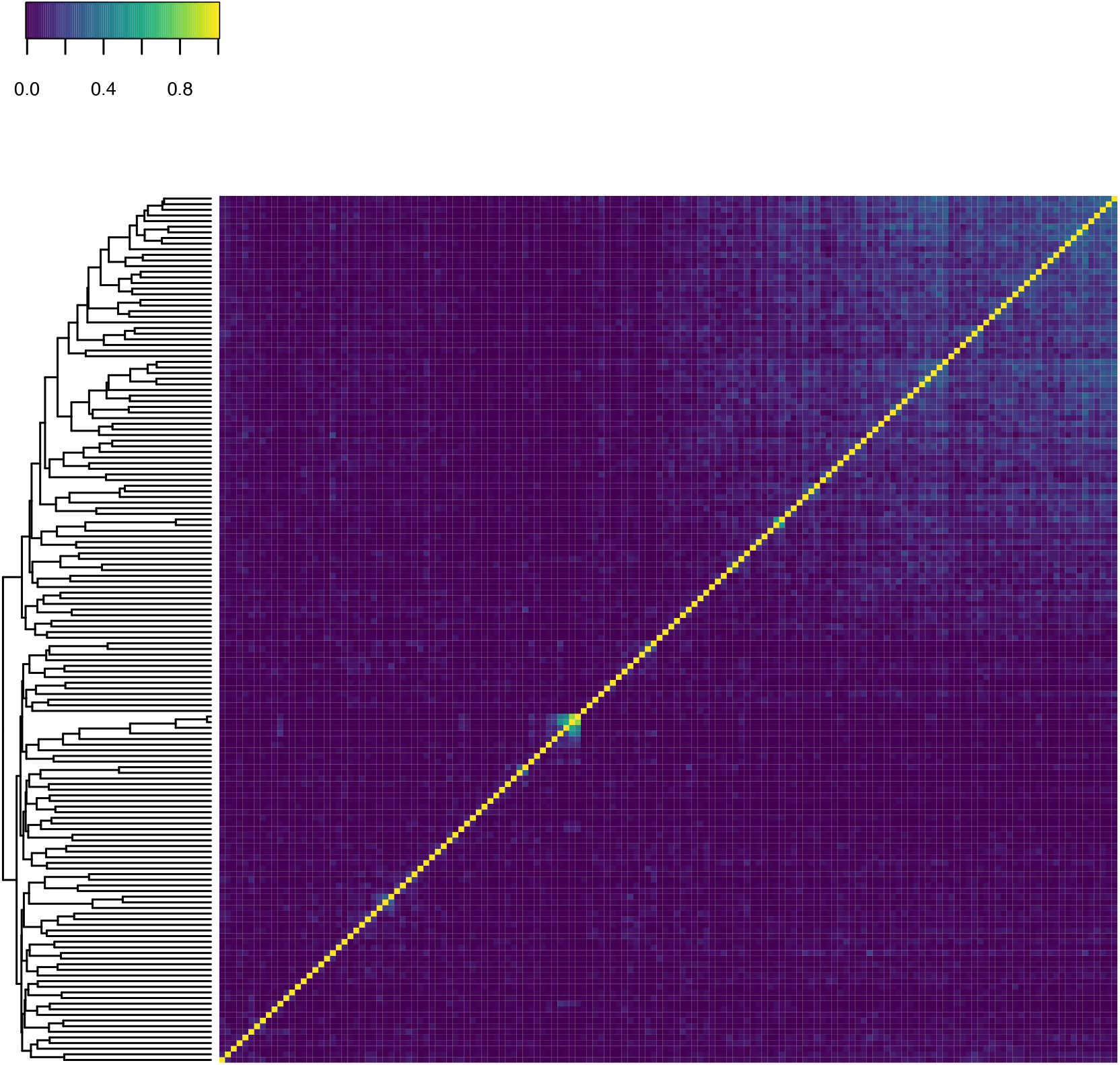
Heatmap of r^2^ values alongside a dendrogram of the 154 agricultural-associated SNPs identified through CMH tests across paired contemporary natural-agricultural samples, illustrating independence among focal LD-clumped CMH outliers.

**Fig S12.**
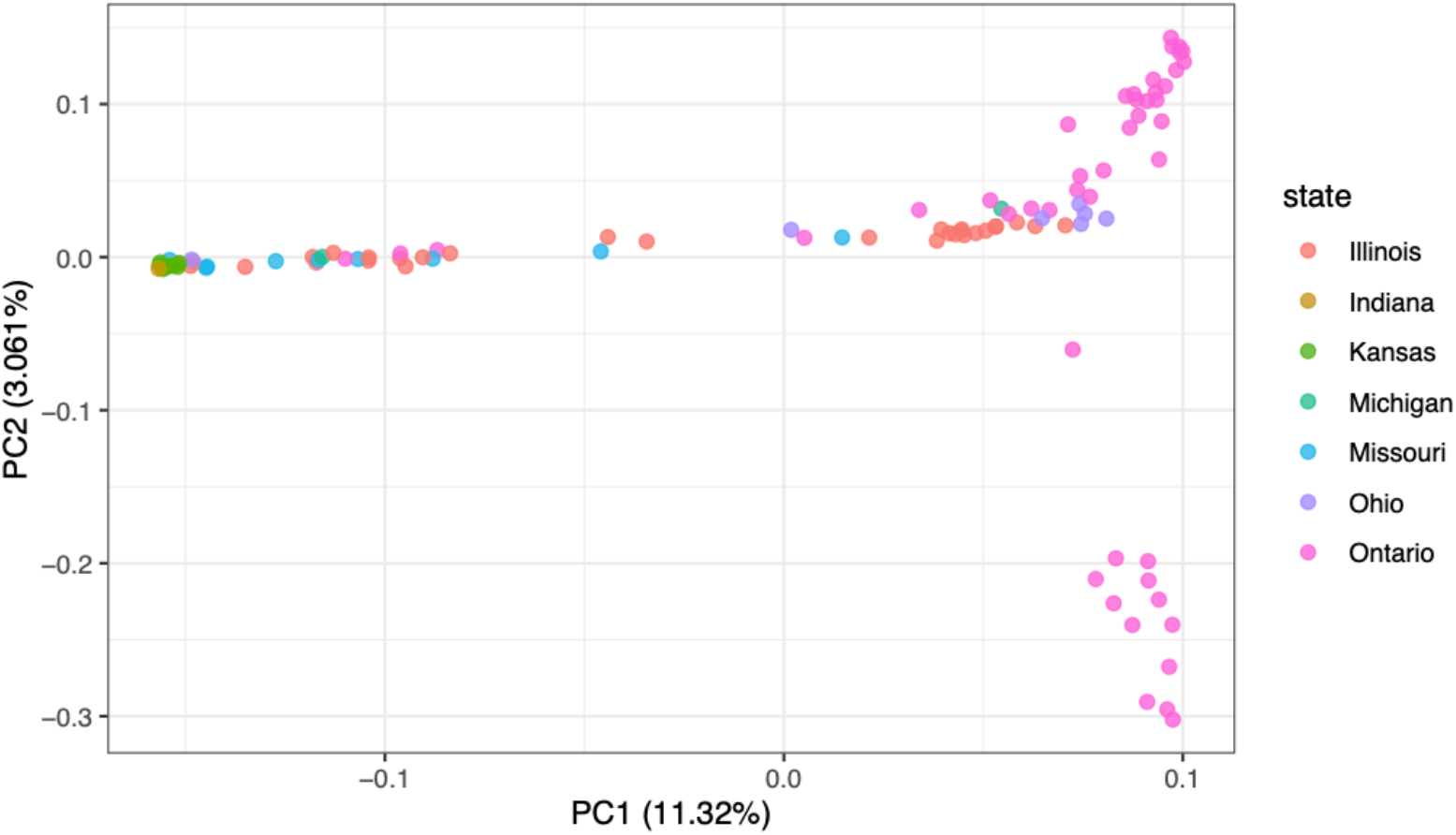
PCA of genome-wide genotypes from herbarium samples, colored by state/province.

**Fig S13.**
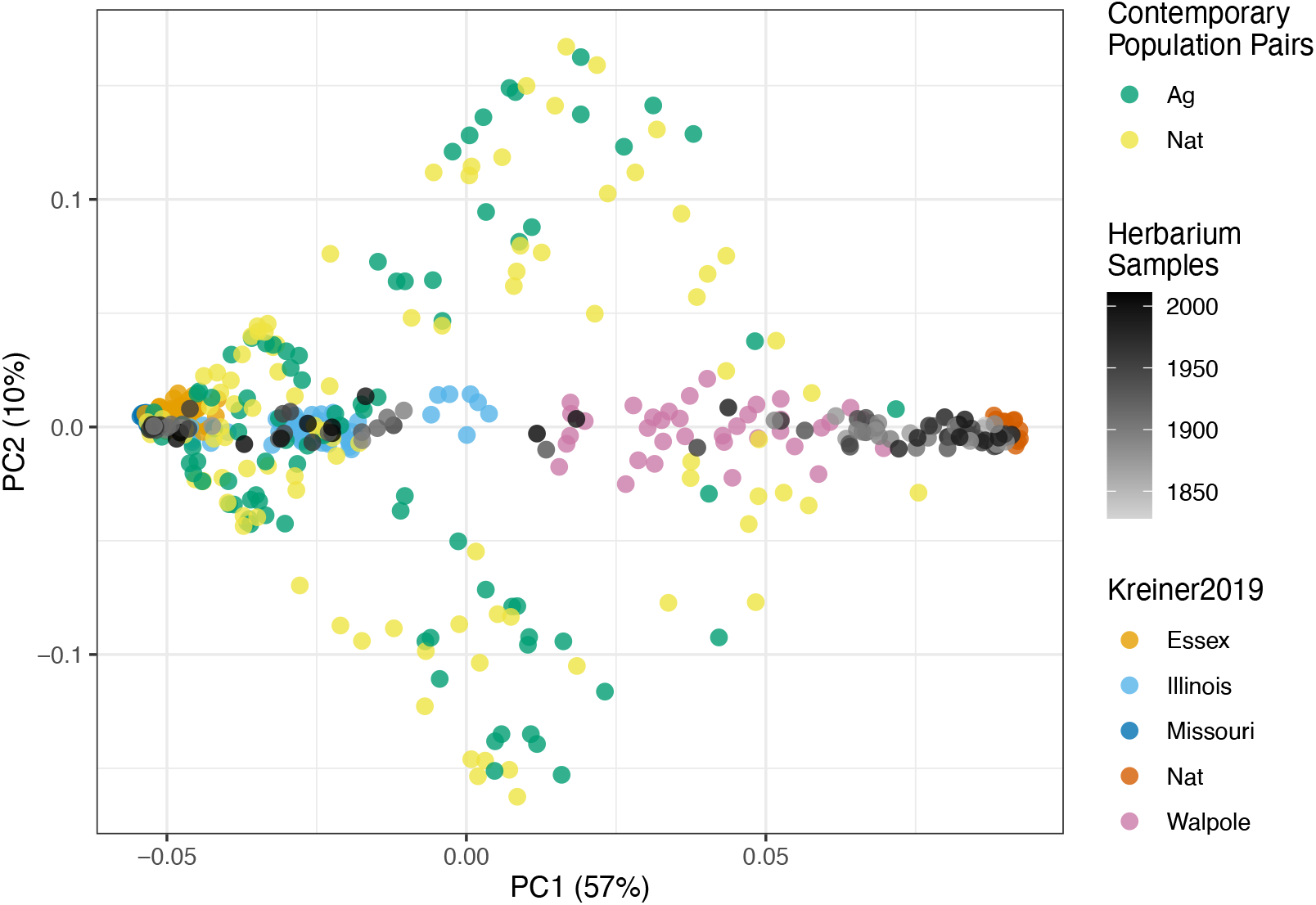
PCA of genome-wide genotypes from 457 *A. tuberculatus* specimens, including 108 herbarium samples, contemporary paired populations (*25*) (n=187), and 21 populations from 5 geographic regions (*47*) (n=162).

**Table S1.**
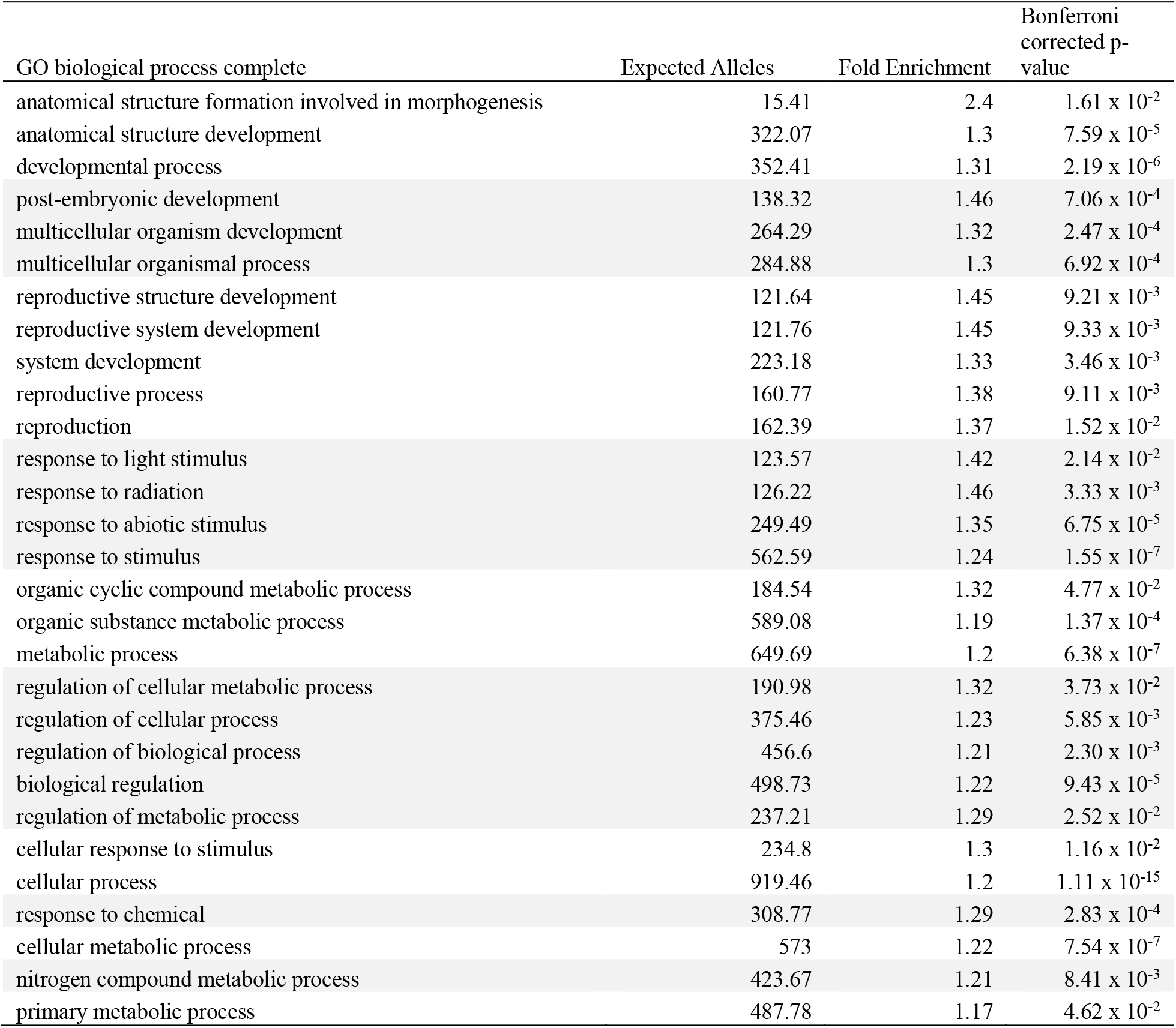
GO Enrichment results for the top 0.1% CMH outliers (n=7264 SNPs, 1650 orthologous genes in *Arabidopsis thaliana*).

**Table S2.** Gene and orthologue information for the 50 SNPs with the most significant CMH p-values, sorted by Scaffold and then CMH p-value. AMATA=*Amaranthus tuberculatus*, AT=*Arabidopsis thaliana*. Blastn top-hit only shown for SNPs with no orthologue identified in our annotation or using orthofinder.

| Scaffold | Position | CMH p-value | AMATA gene | AT gene | Orthologue | Blastn |
| --- | --- | --- | --- | --- | --- | --- |
| 1 | 59264411 | 6.08 x 10<sup>-10</sup> | 2592 | NA | NA | Nuclear Fusion Defective 4-like |
| 1 | 55158523 | 3.29 x 10<sup>-9</sup> | 2285 | At4g34215 | SGNH-hydrolase |  |
| 1 | 54510509 | 8.91 x 10<sup>-9</sup> | 2244 | NA | NA | NA |
| 1 | 12324718 | 1.93 x 10<sup>-7</sup> | 825 | AT5G09550.1 | Guanosine nucleotide diphosphate dissociation inhibitor (GDI) |  |
| 1 | 64789405 | 2.39 x 10<sup>-7</sup> | 3031 | AT4G38380.4 | Protein DETOXIFICATION 452C chloroplastic (DTX45) |  |
| 10 | 15265237 | 7.25 x 10<sup>-8</sup> | 22183 | NA | NA | NA |
| 10 | 26222541 | 1.03 x 10<sup>-7</sup> | 22540 | AT3G29385.1 | Dentin sialophosphoprotein-like protein |  |
| 10 | 5374087 | 2.74 x 10<sup>-7</sup> | 21789 | NA | NA | NA |
| 11 | 24079642 | 8.55 x 10<sup>-11</sup> | 25990 | AT5G63460.1 | Lower temperature 1 |  |
| 11 | 24078348 | 3.09 x 10<sup>-10</sup> | 25990 | AT5G63460.1 | Lower temperature 1 |  |
| 11 | 24086055 | 1.10 x 10<sup>-9</sup> | 25990 | AT5G63460.1 | Lower temperature 1 |  |
| 11 | 24006946 | 2.18 x 10<sup>-9</sup> | 25984 | AT5G14220.4 | PPO2 |  |
| 11 | 24062979 | 2.36 x 10<sup>-9</sup> | 25989 | AT5G50380.1 | Exocyst complex component EXO70B1 |  |
| 11 | 24080739 | 4.33 x 10<sup>-9</sup> | 25990 | AT5G63460.1 | Lower temperature 1 |  |
| 11 | 32783081 | 7.71 x 10<sup>-9</sup> | 26623 | NA | NA | NA |
| 11 | 24048769 | 1.30 x 10<sup>-8</sup> | 25988 | AT4G14110.1 | COP9 signalosome complex subunit 8 (CSN8) |  |
| 11 | 24047019 | 4.13 x 10<sup>-8</sup> | 25988 | AT4G14110.1 | COP9 signalosome complex subunit 8 (CSN8) |  |
| 11 | 24083434 | 4.40 x 10<sup>-8</sup> | 25990 | AT5G63460.1 | Lower temperature 1 |  |
| 11 | 24070607 | 4.79 x 10<sup>-8</sup> | 25989 | AT5G50380.1 | Exocyst complex component EXO70B1 |  |
| 11 | 26024805 | 1.35 x 10<sup>-7</sup> | 26127 | AT1G75125.1 | Plastid transcriptionally active protein |  |
| 11 | 25369807 | 1.47 x 10<sup>-7</sup> | 26088 | AT5G39610.1 | Nucleobase-ascorbate transporter 6 (NAC6) |  |
| 11 | 24021382 | 1.69 x 10<sup>-7</sup> | 25985 | AT5G16550.1 | Ldap interacting protein |  |
| 11 | 24024155 | 1.69 x 10<sup>-7</sup> | 25985 | AT5G16550.2 | Ldap interacting protein |  |
| 11 | 24048722 | 2.07 x 10<sup>-7</sup> | 25988 | AT4G14110.1 | Constitutive photomorphogenic 9 |  |
| 11 | 24046969 | 2.55 x 10<sup>-7</sup> | 25988 | AT4G14110.1 | Constitutive photomorphogenic 9 |  |
| 12 | 29335314 | 7.69 x 10<sup>-10</sup> | 24987 | ATMG00310.1 | Orf154 |  |
| 12 | 29335422 | 3.09 x 10<sup>-9</sup> | 24987 | ATMG00310.1 | Orf154 |  |
| 12 | 29327164 | 3.31 x 10<sup>-9</sup> | 24987 | ATMG00310.1 | Orf154 |  |
| 12 | 29343299 | 1.81 x 10<sup>-8</sup> | 24987 | ATMG00310.1 | Orf154 |  |
| 12 | 29336458 | 3.50 x 10<sup>-8</sup> | 24987 | ATMG00310.1 | Orf154 |  |
| 12 | 29333763 | 5.83 x 10<sup>-8</sup> | 24987 | ATMG00310.1 | Orf154 |  |
| 12 | 11427671 | 7.91 x 10<sup>-8</sup> | 24182 | NA | NA | PPX2L |
| 12 | 29328119 | 1.21 x 10<sup>-7</sup> | 24987 | ATMG00310.1 | Orf154 |  |
| 12 | 11429925 | 2.10 x 10<sup>-7</sup> | 24182 | NA | NA | PPX2L |
| 12 | 29333575 | 2.30 x 10<sup>-7</sup> | 24987 | ATMG00310.1 | Orf154 |  |
| 12 | 29333622 | 2.30 x 10<sup>-7</sup> | 24987 | ATMG00310.2 | Orf155 |  |
| 13 | 35715956 | 8.71 x 10<sup>-9</sup> | 19321 | NA | Calmodulin (Physarum polycephalum OX%253D5791) |  |
| 13 | 38847060 | 6.31 x 10<sup>-8</sup> | 19605 | NA | NA | WIP2-like protein |
| 13 | 26858220 | 8.73 x 10<sup>-8</sup> | 18867 | NA | NA | NA |
| 13 | 33084574 | 1.93 x 10<sup>-7</sup> | 19117 | AT4G14310.2 | KIN14B-interacting protein |  |
| 13 | 41600578 | 2.63 x 10<sup>-7</sup> | 19862 | AT3G24160.1 | Putative type 1 membrane protein |  |
| 2 | 53114550 | 8.86 x 10<sup>-8</sup> | 5599 | NA | NA | NA |
| 2 | 10688517 | 9.38 x 10<sup>-8</sup> | 3997 | AT3G09630.1 | Suppressor of acaulis 56 (sac56) |  |
| 3 | 4797458 | 8.95 x 10<sup>-9</sup> | 6345 | NA | NA | ATHB13 |
| 3 | 1072396 | 9.09 x 10<sup>-9</sup> | 5998 | AT1G23820.1 | Spermidine synthase 1 |  |
| 3 | 1072448 | 8.96 x 10<sup>-8</sup> | 5998 | AT1G23820.1 | Spermidine synthase 1 |  |
| 3 | 16438213 | 1.27 x 10<sup>-7</sup> | 7059 | AT1G10150.1 | Carbohydrate-binding protein |  |
| 3 | 5632229 | 1.80 x 10<sup>-7</sup> | 6414 | AT4G09650.1 | Atp synthase delta-subunit gene (atpd) |  |
| 6 | 18669007 | 1.54 x 10<sup>-7</sup> | 15047 | AT4G35830.1 | Aconitase 1 (ACO1) |  |
| 8 | 15398786 | 1.13 x 10<sup>-8</sup> | 20978 | AT3G14310.1 | Pectin methylesterase 3 |  |

**Table S3.** Selection-migration differentiation statistics for 8 resistance alleles, along with maximum likelihood estimates of selection based on allele frequency through time. Ag, agricultural sites; Nat, natural sites. Cost and benefit estimates shown here for the additive (*h*=0.5) case. ***S***_**1960-2018**_ represents the maximum likelihood estimate of selection from the binomial sampling equation of allele frequency change from 1960-2018 based on a diploid model of selection, along with the associated 95% confidence interval. In the main text, Cost:Ben is only presented for alleles with significant differences in frequency between natural and agricultural habitats (*).

| | Ag freq | Nat freq | Ag benefit ($s_A/m_A$) | Nat cost ($s_N/m_N$) | Cost:Ben | $s_{1960-2018}$ | 95% CI of $s$ |
| --- | --- | --- | --- | --- | --- | --- | --- |
| **PPOdel\*** | 0.334 | 0.0464 | 2.59 | 13.00 | 5.03 | 1.16 | 0.19, ∞ |
| **EPSPSamp\*** | 0.496 | 0.236 | 2.08 | 2.88 | 1.39 | NA | NA |
| **EPSPS106** | 0.127 | 0.087 | 0.72 | 1.01 | 1.40 | 1.12 | 0.11, ∞ |
| **ALS122** | 0.0301 | 0 | 2.06 | NA | NA | -0.01 | $-\infty$, ∞ |
| **ALS197** | 0.0132 | 0.0169 | -0.57 | -0.45 | 1.26 | 0.29 | $-\infty$, ∞ |
| **ALS574\*** | 0.337 | 0.191 | 1.31 | 1.89 | 1.45 | 0.09 | 0.088, 0.334 |
| **ALS376** | 0.0824 | 0.0358 | 1.23 | 2.70 | 2.19 | 0.59 | 0.046, ∞ |
| **ALS653** | 0.0596 | 0.0627 | -0.11 | -0.11 | 1 | 0.55 | 0.058, ∞ |

**Table S4.** The top 15 loci with the strongest estimates of selection (*s*) between 1970 and 2018 based on logistic regression. AT = *Arabidopsis thaliana*

| *s* | p-value<br>of *s* | SE<br>of *s* | Allele freq<br>change | Scaffold | position | AMATA annotated gene | AT orthologue | AT gene<br>name |
| --- | --- | --- | --- | --- | --- | --- | --- | --- |
| 0.106 | $4.7 \times 10^{-4}$ | 0.030 | 0.846 | Scaffold_11 | 26068182 | AMATA_chromosomes_26131 | NA | NA* |
| 0.081 | $1.1 \times 10^{-4}$ | 0.021 | 0.823 | Scaffold_2 | 10755264 | AMATA_chromosomes_03999 | Subtilase family protein | AT5G58840 |
| 0.057 | $2.4 \times 10^{-3}$ | 0.019 | 0.512 | Scaffold_10 | 22995135 | AMATA_chromosomes_22407 | phosphotyrosyl phosphatase activator (PTPA family protein) | AT4G08960 |
| 0.052 | $8.2 \times 10^{-3}$ | 0.020 | 0.402 | Scaffold_11 | 26024805 | AMATA_chromosomes_26127 | plastid transcriptionally active protein | AT1G75125* |
| 0.052 | $6.1 \times 10^{-5}$ | 0.013 | 0.646 | Scaffold_3 | 14213167 | AMATA_chromosomes_06976 | WRKY DNA-binding protein 13 | AT4G39410 |
| 0.047 | $2.5 \times 10^{-7}$ | 0.009 | 0.783 | Scaffold_10 | 36863790 | AMATA_chromosomes_23250 | hypothetical protein | AT1G36320 |
| 0.046 | $1.3 \times 10^{-1}$ | 0.030 | 0.162 | Scaffold_3 | 49332690 | AMATA_chromosomes_08117 | NA | NA |
| 0.045 | $2.9 \times 10^{-5}$ | 0.011 | 0.677 | Scaffold_6 | 18669007 | AMATA_chromosomes_15047 | ACO1 | AT4G35830 |
| 0.043 | $8.6 \times 10^{-5}$ | 0.011 | 0.599 | Scaffold_12 | 21792253 | AMATA_chromosomes_24760 | NA | NA |
| 0.043 | $4.6 \times 10^{-8}$ | 0.008 | 0.888 | Scaffold_3 | 5632229 | AMATA_chromosomes_06414 | ATPD (F-type H<sup>+</sup>-transporting ATPase subunit delta) | AT4G09650 |
| 0.036 | $8.1 \times 10^{-6}$ | 0.008 | 0.666 | Scaffold_12 | 5658420 | AMATA_chromosomes_23853 | NA | NA |
| 0.033 | $4.7 \times 10^{-7}$ | 0.007 | 0.831 | Scaffold_10 | 20690917 | AMATA_chromosomes_22321 | CCB2 (chaperone DUF2930) | AT5G52110 |
| 0.032 | $1.8 \times 10^{-7}$ | 0.006 | 0.819 | Scaffold_10 | 16005835 | AMATA_chromosomes_22202 | NA | NA |
| 0.032 | $3.9 \times 10^{-5}$ | 0.008 | 0.708 | Scaffold_10 | 24312710 | AMATA_chromosomes_22452 | BPA4 (RNA-binding RRM/RBD/RNP motifs family protein, AT1G14340) | AT1G14340 |
| 0.031 | $5.2 \times 10^{-6}$ | 0.007 | 0.764 | Scaffold_2 | 17891465 | AMATA_chromosomes_04209 | NA | NA |

\* ~2 Mb from PPO

## References and Notes

1. K. Klein Goldewijk, A. Beusen, J. Doelman, E. Stehfest, Anthropogenic land use estimates for the Holocene – HYDE 3.2. Earth Syst. Sci. Data. 9, 927–953 (2017).

2. G. F. Sassenrath, P. Heilman, E. Luschei, G. L. Bennett, G. Fitzgerald, P. Klesius, W. Tracy, J. R. Williford, P. V. Zimba, Technology, complexity and change in agricultural production systems. Renew. Agric. Food Syst. 23, 285–295 (2008).

3. T. E. Crews, W. Carton, L. Olsson, Is the future of agriculture perennial? Imperatives and opportunities to reinvent agriculture by shifting from annual monocultures to perennial polycultures. Glob. sustain. 1 (2018), doi:10.1017/sus.2018.11.

4. E. Malaj, L. Freistadt, C. A. Morrissey, Spatio-temporal patterns of crops and agrochemicals in Canada over 35 years. Front. Environ. Sci. 8 (2020), doi:10.3389/fenvs.2020.556452.

5. J. Fernandez-Cornejo, R. F. Nehring, C. Osteen, S. Wechsler, A. Martin, A. Vialou, Pesticide use in U.s. agriculture: 21 selected crops, 1960-2008. SSRN Electron. J. (2014), doi:10.2139/ssrn.2502986.

6. N. E. Borlaug, Contributions of conventional plant breeding to food production. Science. 219, 689–693 (1983).

7. H. J. Beckie, K. N. Harker, L. M. Hall, S. I. Warwick, A. Légère, P. H. Sikkema, G. W. Clayton, A. G. Thomas, J. Y. Leeson, G. Séguin-Swartz, M.-J. Simard, A decade of herbicide-resistant crops in Canada. Can. J. Plant Sci. 86, 1243–1264 (2006).

8. C. Mann, Reseeding the Green Revolution. Science (1997), doi:10.1126/science.277.5329.1038.

9. P. L. Pingali, Green revolution: impacts, limits, and the path ahead. Proc. Natl. Acad. Sci. U. S. A. 109, 12302–12308 (2012).

10. P. Pellegrini, R. J. Fernández, Crop intensification, land use, and on-farm energy-use efficiency during the worldwide spread of the green revolution. Proc. Natl. Acad. Sci. U. S. A. 115, 2335–2340 (2018).

11. J. Mallet, The evolution of insecticide resistance: Have the insects won? Trends Ecol. Evol. 4, 336–340 (1989).

12. C. Délye, M. Jasieniuk, V. Le Corre, Deciphering the evolution of herbicide resistance in weeds. Trends Genet. 29, 649–658 (2013).

13. N. J. Hawkins, C. Bass, A. Dixon, P. Neve, The evolutionary origins of pesticide resistance. Biol. Rev. Camb. Philos. Soc. (2018), doi:10.1111/brv.12440.

14. F. Gould, Z. S. Brown, J. Kuzma, Wicked evolution: Can we address the sociobiological dilemma of pesticide resistance? Science. 360, 728–732 (2018).

15. F. Zabel, R. Delzeit, J. M. Schneider, R. Seppelt, W. Mauser, T. Václavík, Global impacts of future cropland expansion and intensification on agricultural markets and biodiversity. Nat. Commun. 10, 2844 (2019).

16. J. Sauer, REVISION OF THE DIOECIOUS AMARANTHS. Madroño. 13, 5–46 (1955).

17. K. E. Waselkov, K. M. Olsen, Population genetics and origin of the native North American agricultural weed waterhemp (Amaranthus tuberculatus; Amaranthaceae). Am. J. Bot. 101, 1726–1736 (2014).

18. M. Costea, S. E. Weaver, F. J. Tardif, The Biology of Invasive Alien Plants in Canada. 3. Amaranthus tuberculatus (Moq.) Sauer var. rudis (Sauer) Costea & Tardif. Can. J. Plant Sci. 85, 507–522 (2005).

19. P. J. Tranel, Herbicide resistance in Amaranthus tuberculatus†. Pest Manag. Sci. 77, 43–54 (2021).

20. J. Hermisson, P. S. Pennings, Soft sweeps: molecular population genetics of adaptation from standing genetic variation. Genetics. 169, 2335–2352 (2005).

21. R. D. H. Barrett, D. Schluter, Adaptation from standing genetic variation. Trends Ecol. Evol. 23, 38–44 (2008/1).

22. M. Alleaume-Benharira, I. R. Pen, O. Ronce, Geographical patterns of adaptation within a species’ range: interactions between drift and gene flow. J. Evol. Biol. 19, 203–215 (2006).

23. R. I. Colautti, S. C. H. Barrett, Rapid adaptation to climate facilitates range expansion of an invasive plant. Science. 342, 364–366 (2013).

24. M. Szucs, M. L. Vahsen, B. A. Melbourne, C. Hoover, C. Weiss-Lehman, R. A. Hufbauer, Rapid adaptive evolution in novel environments acts as an architect of population range expansion. Proc. Natl. Acad. Sci. U. S. A. 114, 13501–13506 (2017).

25. J. M. Kreiner, A. Caballero, S. I. Wright, J. R. Stinchcombe, Selective ancestral sorting and de novo evolution in the agricultural invasion of Amaranthus tuberculatus. Evolution (2021), doi:10.1111/evo.14404.

26. F. E. Dayan, S. O. Duke, “Chapter 81 - Protoporphyrinogen Oxidase-Inhibiting Herbicides” in Hayes’ Handbook of Pesticide Toxicology, R. Krieger, Ed. (Academic Press, New York, 2010), pp. 1733–1751.

27. J. G. Moraes, T. R. Butts, V. M. Anunciato, J. D. Luck, W. C. Hoffmann, U. R. Antuniassi, G. R. Kruger, Nozzle selection and adjuvant impact on the efficacy of glyphosate and PPO-inhibiting herbicide tank-mixtures. Agronomy (Basel). 11, 754 (2021).

28. W. Moeder, O. Del Pozo, D. A. Navarre, G. B. Martin, D. F. Klessig, Aconitase plays a role in regulating resistance to oxidative stress and cell death in Arabidopsis and Nicotiana benthamiana. Plant Mol. Biol. 63, 273–287 (2007).

29. P. A. Ribone, M. Capella, R. L. Chan, Functional characterization of the homeodomain leucine zipper I transcription factor AtHB13 reveals a crucial role in Arabidopsis development. J. Exp. Bot. 66, 5929–5943 (2015).

30. J. V. Cabello, R. L. Chan, The homologous homeodomain-leucine zipper transcription factors HaHB1 and AtHB13 confer tolerance to drought and salinity stresses via the induction of proteins that stabilize membranes. Plant Biotechnol. J. 10, 815–825 (2012).

31. S. Guénin, J. Hardouin, F. Paynel, K. Müller, G. Mongelard, A. Driouich, P. Lerouge, A. R. Kermode, A. Lehner, J.-C. Mollet, J. Pelloux, L. Gutierrez, A. Mareck, AtPME3, a ubiquitous cell wall pectin methylesterase of Arabidopsis thaliana, alters the metabolism of cruciferin seed storage proteins during post-germinative growth of seedlings. J. Exp. Bot. 68, 1083–1095 (2017).

32. S. Zhou, L. Jia, H. Chu, D. Wu, X. Peng, X. Liu, J. Zhang, J. Zhao, K. Chen, L. Zhao, Arabidopsis CaM1 and CaM4 Promote Nitric Oxide Production and Salt Resistance by Inhibiting S-Nitrosoglutathione Reductase via Direct Binding. PLoS Genet. 12, e1006255 (2016).

33. C. Dai, Y. Lee, I. C. Lee, H. G. Nam, J. M. Kwak, Calmodulin 1 Regulates Senescence and ABA Response in Arabidopsis. Front. Plant Sci. 9, 803 (2018).

34. H. Guo, H. Yang, T. C. Mockler, C. Lin, Regulation of flowering time by Arabidopsis photoreceptors. Science. 279, 1360–1363 (1998).

35. T. Mockler, H. Yang, X. Yu, D. Parikh, Y.-C. Cheng, S. Dolan, C. Lin, Regulation of photoperiodic flowering by Arabidopsis photoreceptors. Proc. Natl. Acad. Sci. U. S. A. 100, 2140–2145 (2003).

36. W. Wang, X. Lu, L. Li, H. Lian, Z. Mao, P. Xu, T. Guo, F. Xu, S. Du, X. Cao, S. Wang, H. Shen, H.-Q. Yang, Photoexcited CRYPTOCHROME1 Interacts with Dephosphorylated BES1 to Regulate Brassinosteroid Signaling and Photomorphogenesis in Arabidopsis. Plant Cell. 30, 1989–2005 (2018).

37. J. Li, Y. Li, S. Chen, L. An, Involvement of brassinosteroid signals in the floral-induction network of Arabidopsis. J. Exp. Bot. 61, 4221–4230 (2010).

38. F. E. Dayan, P. R. Daga, S. O. Duke, R. M. Lee, P. J. Tranel, R. J. Doerksen, Biochemical and structural consequences of a glycine deletion in the α-8 helix of protoporphyrinogen oxidase. Biochimica et Biophysica Acta (BBA) - Proteins and Proteomics. 1804, 1548–1556 (2010).

39. H. M. Cockerton, S. S. Kaundun, L. Nguyen, S. J. Hutchings, R. P. Dale, A. Howell, P. Neve, Fitness cost associated with enhanced EPSPS gene copy number and glyphosate resistance in an Amaranthus tuberculatus population. Cold Spring Harbor Laboratory (2021), p. 2021.01.09.426028,, doi:10.1101/2021.01.09.426028.

40. M. M. Vila-Aiub, Fitness of Herbicide-Resistant Weeds: Current Knowledge and Implications for Management. Plants. 8 (2019), doi:10.3390/plants8110469.

41. Kreiner, J.M. Latorre, S.M., Burbano, H.A., Stinchcombe, J.R., Otto, S.P., Weigel, D., Wright, S., Materials and Methods for “200 years of agricultural adaptation and range expansion in a native weed.” Science.

42. N. H. Barton, The “New Synthesis.” Proceedings of the National Academy of Sciences. 119, e2122147119 (2022).

43. M. Przeworski, The signature of positive selection at randomly chosen loci. Genetics. 160, 1179–1189 (2002).

44. Y. Kim, R. Nielsen, Linkage disequilibrium as a signature of selective sweeps. Genetics. 167, 1513–1524 (2004).

45. J. M. Kreiner, G. Sandler, A. J. Stern, P. J. Tranel, D. Weigel, J. Stinchcombe, S. I. Wright, Repeated origins, widespread gene flow, and allelic interactions of target-site herbicide resistance mutations. Elife. 11, e70242 (2022).

46. J. Sauer, Recent Migration and Evolution of the Dioecious Amaranths. Evolution. 11, 11–31 (1957).

47. J. M. Kreiner, D. A. Giacomini, F. Bemm, B. Waithaka, J. Regalado, C. Lanz, J. Hildebrandt, P. H. Sikkema, P. J. Tranel, D. Weigel, J. R. Stinchcombe, S. I. Wright, Multiple modes of convergent adaptation in the spread of glyphosate-resistant Amaranthus tuberculatus. Proc. Natl. Acad. Sci. U. S. A. 116, 21076–21084 (2019).

48. A. Raj, M. Stephens, J. K. Pritchard, fastSTRUCTURE: variational inference of population structure in large SNP data sets. Genetics. 197, 573–589 (2014).

49. M. J. Foes, L. Liu, P. J. Tranel, L. M. Wax, E. W. Stoller, A biotype of common waterhemp (Amaranthus rudis) resistant to triazine and ALS herbicides. Weed Sci. 46, 514–520 (1998).

50. P. J. Tranel, C. W. Riggins, M. S. Bell, A. G. Hager, Herbicide resistances in Amaranthus tuberculatus: a call for new options. J. Agric. Food Chem. 59, 5808–5812 (2011).

51. Q. C. B. Cronk, J. L. Fuller, Plant invaders: The threat to natural ecosystems (Routledge, 2014).

52. M. Exposito-Alonso, 500 Genomes Field Experiment Team, H. A. Burbano, O. Bossdorf, R. Nielsen, D. Weigel, Natural selection on the Arabidopsis thaliana genome in present and future climates. Nature (2019), doi:10.1038/s41586-019-1520-9.

53. J. M. Kreiner, jkreinz/TemporalAdaptation: Oct92022_TemporalAdaptationCode, version 1.0.0 (2022),, doi:10.5281/zenodo.7178764.

## References for Methods & Materials

54. S. M. Latorre, P. L. M. Lang, H. A. Burbano, R. M. Gutaker, Isolation, Library Preparation, and Bioinformatic Analysis of Historical and Ancient Plant DNA. Curr Protoc Plant Biol. 5, e20121 (2020).

55. S. Chen, Y. Zhou, Y. Chen, J. Gu, fastp: an ultra-fast all-in-one FASTQ preprocessor. Bioinformatics. 34 (2018), pp. i884–i890.

56. H. Li, R. Durbin, Fast and accurate short read alignment with Burrows-Wheeler transform. Bioinformatics. 25, 1754–1760 (2009).

57. A. Peltzer, G. Jäger, A. Herbig, A. Seitz, C. Kniep, J. Krause, K. Nieselt, EAGER: efficient ancient genome reconstruction. Genome Biol. 17, 60 (2016).

58. A. Ginolhac, M. Rasmussen, M. T. P. Gilbert, E. Willerslev, L. Orlando, mapDamage: testing for damage patterns in ancient DNA sequences. Bioinformatics. 27, 2153–2155 (2011).

59. S. Purcell, B. Neale, K. Todd-Brown, L. Thomas, M. A. R. Ferreira, D. Bender, J. Maller, P. Sklar, P. I. W. de Bakker, M. J. Daly, P. C. Sham, PLINK: a tool set for whole-genome association and population-based linkage analyses. Am. J. Hum. Genet. 81, 559–575 (2007).

60. D. M. Emms, S. Kelly, OrthoFinder: solving fundamental biases in whole genome comparisons dramatically improves orthogroup inference accuracy. Genome Biol. 16, 157 (2015).

61. S. F. Altschul, W. Gish, W. Miller, E. W. Myers, D. J. Lipman, Basic local alignment search tool. J. Mol. Biol. 215, 403–410 (1990).

62. Z. A. Szpiech, R. D. Hernandez, selscan: an efficient multithreaded program to perform EHH-based scans for positive selection. Mol. Biol. Evol. 31, 2824–2827 (2014).

63. O. Delaneau, J.-F. Zagury, J. Marchini, Improved whole-chromosome phasing for disease and population genetic studies. Nat. Methods. 10, 5–6 (2013).

64. F. J. Tardif, I. Rajcan, M. Costea, A mutation in the herbicide target site acetohydroxyacid synthase produces morphological and structural alterations and reduces fitness in Amaranthus powellii. New Phytol. 169, 251–264 (2006).

65. M. M. Vila-Aiub, S. S. Goh, T. A. Gaines, H. Han, R. Busi, Q. Yu, S. B. Powles, No fitness cost of glyphosate resistance endowed by massive EPSPS gene amplification in Amaranthus palmeri. Planta. 239, 793–801 (2014).

66. C. Wu, A. S. Davis, P. J. Tranel, Limited fitness costs of herbicide-resistance traits in Amaranthus tuberculatus facilitate resistance evolution. Pest Manag. Sci. 74, 293–301 (2018).

67. S. Gupta, A. Harkess, A. Soble, M. Van Etten, J. Leebens-Mack, R. S. Baucom, Inter-chromosomal linkage disequilibrium and linked fitness cost loci influence the evolution of nontarget site herbicide resistance in an agricultural weed. bioRxiv (2021), p. 2021.04.04.438381,, doi:10.1101/2021.04.04.438381.

68. B. Pasaniuc, S. Sankararaman, G. Kimmel, E. Halperin, Inference of locus-specific ancestry in closely related populations. Bioinformatics. 25, i213–21 (2009).

